# A glutamine-based single ɑ-helix scaffold to target globular proteins

**DOI:** 10.1101/2022.05.06.490931

**Authors:** A. Escobedo, J. Piccirillo, J. Aranda, T. Diercks, B. Topal, M. Biesaga, L. Staby, B. B. Kragelund, J. García, O. Millet, M. Orozco, M. Coles, R. Crehuet, X. Salvatella

**Affiliations:** Institute for Research in Biomedicine (IRB Barcelona), The Barcelona Institute of Science and Technology, Baldiri Reixac 10, 08028 Barcelona, Spain; CIC bioGUNE, Bizkaia, Science and Technology Park bld 801A, 48160 Derio, Bizkaia, Spain; REPIN and Structural Biology and NMR Laboratory, The Linderstrøm-Lang Centre for Protein Science, Department of Biology, Ole Maaloes Vej 5, University of Copenhagen, DK-2200 Copenhagen N, Denmark; Department of Biochemistry and Biomedicine, University of Barcelona, Avinguda Diagonal 645, 08028 Barcelona, Spain; Department of Protein Evolution, Max Planck Institute for Developmental Biology, Max-Planck-Ring 5, 72076 Tubingen, Germany; CSIC-Institute for Advanced Chemistry of Catalonia (IQAC), Jordi Girona 18-26, 08034 Barcelona, Spain; ICREA, Passeig Lluís Companys 23, 08010, Barcelona, Spain

## Abstract

The binding of intrinsically disordered proteins to globular ones often requires the folding of motifs into ɑ-helices. These interactions offer opportunities for therapeutic intervention but their modulation with small molecules is challenging because they bury large surfaces. Linear peptides that display the residues that are key for binding can be targeted to globular proteins when they form stable helices, which in most cases requires their chemical modification. Here we present rules to design peptides that fold into single ɑ-helices by instead concatenating glutamine side chain to main chain hydrogen bonds recently discovered in polyglutamine helices. The resulting peptides are uncharged, contain only natural amino acids, and their sequences can be optimized to interact with specific targets. Our results provide design rules to obtain single ɑ-helices for a wide range of applications in protein engineering and drug design.

## Introduction

Proteins are central to biology as they carry out a wide range of essential functions, from gene regulation to enzymatic catalysis, where their ability to specifically interact with other biomolecules is key. In pharmacology, inhibiting their interactions using drug-like small molecules is a common approach to modulate biological functions relevant to disease. In cases where the binding partner is another protein, the binding interfaces are usually flat and extended^1^, making it challenging to inhibit the interaction with small molecules^2^, and it is generally preferable to target them with antibodies. Yet, despite recent remarkable progress in intracellular antibody delivery^3^, their clinical applications have been limited to targeting extracellular proteins, highlighting the need to develop new molecular tools to inhibit protein-protein interactions.

Peptides have some of the advantages of small molecules, such as their ease of synthesis and rather high bioavailability, and some of the advantages of antibodies, such as their relatively large size. Peptides, therefore, have at least in principle great potential as modulators of protein-protein interactions for pharmacological applications^4,5^. Protein-protein interactions where one partner adopts a helical conformation upon binding are particularly common and especially amenable to inhibition by peptides: an excised linear peptide comprising a suitable sequence can, in principle, inhibit the interaction if it can bind to its partner with high affinity^6^. Linear peptides have a low propensity to fold into stable ɑ-helices, however, and the entropic cost of folding decreases both their affinity for their targets and their stability against proteolytic degradation, highlighting the need to develop new tools to stabilize their helical conformation.

Introducing non-natural amino acids with reactive side chains and connecting them via covalent bonds, as in peptide stapling^7,8^, is a powerful method to constrain linear peptides into conformations akin to ɑ-helices. Stapling is based on the pairwise use of synthetic ɑ-methyl, ɑ-alkenyl amino acids at relative positions *i,i+3*, *i,i+4* or *i,i+7* where their side chains can react by ring closing metathesis when the peptide folds into an ɑ-helix, thus greatly stabilizing this conformation; in some cases, two such connections were concatenated to yield particularly long and stable helices^9^. While such stapled ɑ-helical peptides have shown broad applicability to inhibit protein-protein interactions^10^, they have drawbacks that limit their range of applicability, such as their costly synthesis, limited solubility, and high rigidity.

We recently reported that Gln side chain to main chain hydrogen bonds stabilize the polyglutamine (polyQ) helix, a helical secondary structure formed by the polyQ tract of the androgen receptor (AR)^11,12^. In this structure both the main and side chain amide groups of Gln residues at position *i* donate hydrogens to the main chain CO group of the residues at position *i-4.* The strength of this bifurcate interaction depends on the residue at position *i-4*, with Leu performing particularly well. This interaction has also been observed in the polyQ tract of the protein huntingtin, indicating that it is not specific to AR and can therefore be used in design^13^. As the relative donor and acceptor positions of such interactions are equivalent to those of the covalently linked residues in stapled peptides, we reasoned that they would stabilize the ɑ-helical conformation of linear peptides to enhance their interaction with globular proteins.

To address this hypothesis we design a series of linear, uncharged, and highly soluble peptides and confirm their cooperative folding into single ɑ-helices under physiological conditions. To evaluate the versatility and range of applicability of the design rules we analyze their tolerance towards changes of the acceptor residue, with bulky hydrophobic residues performing best, and introduce a pH-dependent conformational switch; in addition we explore how such bifurcated hydrogen bonds can be combined with electrostatic interactions between side chains to further stabilize helical structure. Most importantly, we show that the sequences of such peptides can be tailored to interact with specific globular proteins. In summary, the simple design rules that we propose can be used to engineer a new class of linear, cooperatively folded helical peptides for use as templates in applications in pharmacology, materials science, synthetic biology and, more generally, in bioengineering.

## Results

### Design of Gln-based single ɑ-helices

Structure prediction algorithms are powerful tools for protein design. Machine learning based algorithms predict the structures of globular proteins^16^ but their applicability to proteins devoid of tertiary interactions is still relatively unexplored^17^. This is illustrated by the results reported for intrinsically disordered (ID) domains possessing polyglutamine (polyQ) tracts in the AlphaFold protein structure database. These tracts are predicted to fold into α-helices^17,18^ whereas experiments indicate that their helical propensity strongly depends on sequence context^11,13,19–22^.The helical propensity of peptides and ID proteins can instead be predicted by algorithms, such as Agadir, that compute the energetics of folding by considering residue helical propensities, capping effects and interactions between side chains^23^. Agadir does not account for the Gln side chain to main chain interactions stabilizing polyQ helices and therefore underestimates the helicity of the polyQ tract in AR: the peptide L_4_Q_16_, excised from AR, has 38% helical propensity according to NMR experiments while Agadir predicts only 3%^11^. To address this we introduced an additional energy term accounting for this interaction (E^L^_i,i+4_) and by minimizing the prediction error (RMSD_Hel_) obtained E^L^_i,i+4_ = -0.6 kcal mol^-1^ for L_4_Q_16_, in the range expected for one hydrogen bond in water^24^. Peptides with a short polyQ tract did not require this correction term (Fig. S1a), however, and E^L^_i,i+4_ depended on tract length: it increased from -0.4 kcal mol^-1^ for L_4_Q_8_ to -0.7 kcal mol^-1^ for L_4_Q_20_. The effective strength of the Gln_i+4_→Leu_i_ interactions in AR peptides depends thus on the number of equivalent interactions following them in the sequence, indicating cooperativity.

To analyze the origin of this behavior and exploit it for peptide design, we studied four peptides of identical amino acid composition but with two potential Gln_i+4_→Leu_i_ interactions (pink arrows in Fig. 1a) at different relative positions. The first such interaction is common to all peptides and the second one is shifted 1 (peptide P1-5), 2 (P2-6), 3 (P3-7) or 5 (P5-9) positions towards the C-terminus. After confirming that they were monomeric under our experimental conditions by size exclusion chromatography coupled to multiple angle light scattering (SEC-MALS) (Fig. S1d), we used solution state nuclear magnetic resonance (NMR) spectroscopy to probe their conformation by exploiting the dependence of ^13^C_ɑ_ and ^1^H_ɑ_ NMR chemical shifts on secondary structure, where larger ^13^C_ɑ_ and lower ^1^H_ɑ_ shifts indicate higher helicity^25^. The NMR spectra showed P3-7 is the most helical peptide (Fig. 1a), indicating that the strength of two Gln_i+4_→Leu_i_ interactions is maximal when the donor of the first (Gln11 in P3-7) and the acceptor of the second (Leu10) interaction share a peptide bond such that the two interactions are concatenated; these results were confirmed by circular dichroism (CD) spectroscopy (Fig. S1c).

**Fig. 1.**
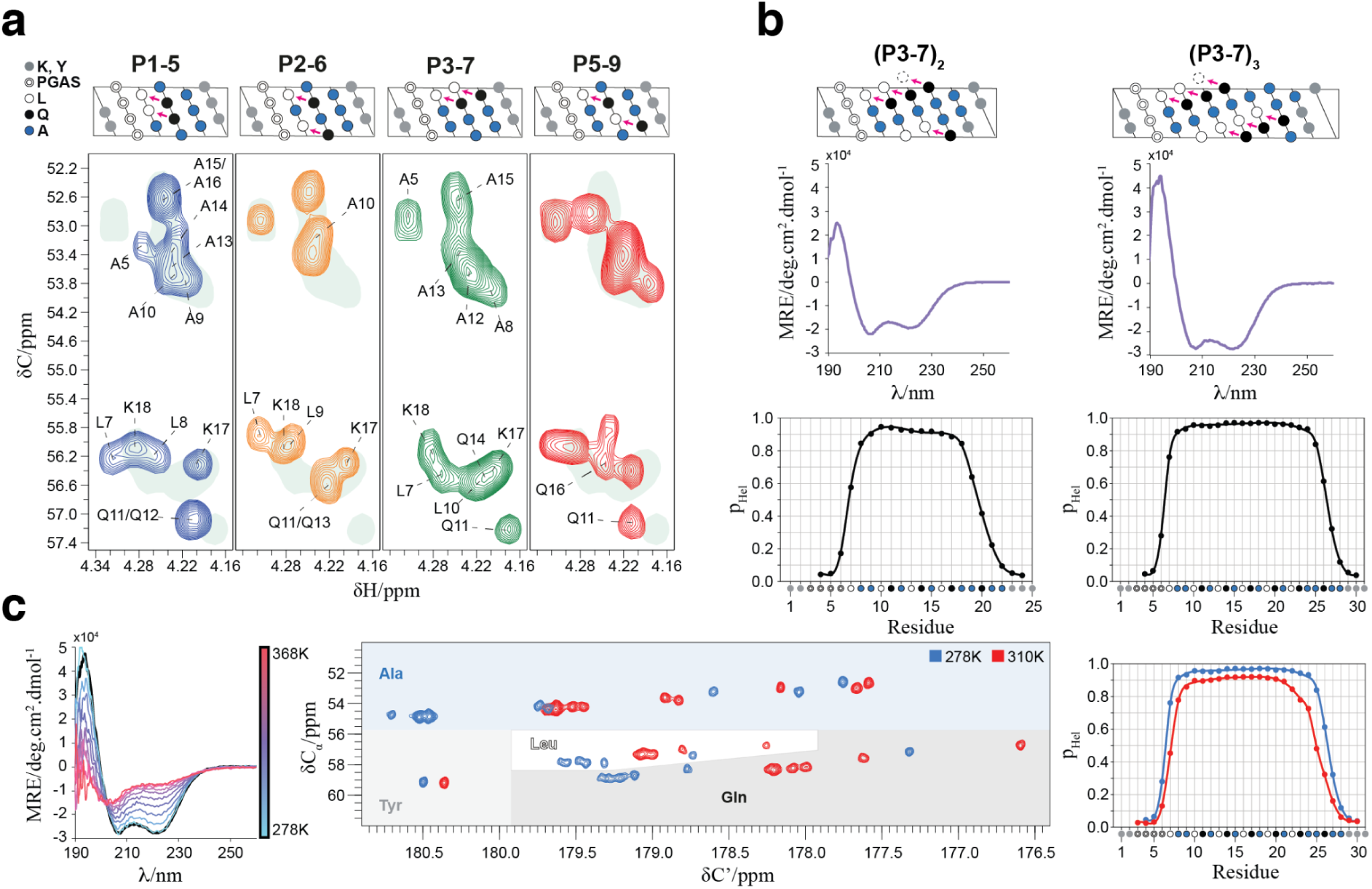
Design of single ɑ-helices stabilized by Gln side chain to main chain hydrogen bonds. **a** Top: representation, as helical projections, of the peptides used to investigate cooperativity, where pink arrows indicate putative side chain to main chain interactions. Bottom: selected region of the 2D ^1^H,^13^C HSQC spectra of the peptides at 278 K, where the shaded area corresponds to that of peptide P3-7. **b** Top: representation, as helical projections, of peptides (P3-7)_2_ and (P3-7)_3_, as in panel a. Middle: CD spectra at 278 K. Bottom: residue-specific helical propensities as obtained from the C’, C_α_, N^H^, and H^N^ NMR chemical shifts using CheSPI^14,15^. **c** Left: CD spectra of peptides (P3-7)_2_ and (P3-7)_3_ obtained at temperatures between 278 K (blue) and 368 K (red) in steps of 10 K; the spectrum obtained at 278 K after refolding is shown in black. Center: superposition of a selected region of the ^13^C detected 2D CACO NMR spectra of peptide (P3-7)_3_ recorded at 278 K (blue) and 310 K (red), with an indication of amino acid specific regions as shades. Right: residue-specific helical propensities as obtained from the NMR chemical shifts at 278 and 310 K as in **b**.

We then used CD spectroscopy to study the secondary structure of peptides containing two and three pairs of concatenated Gln_i+4_→Leu_i_ interactions, (P3-7)_2_ and (P3-7)_3_, and obtained that they are highly helical and monomeric (Θ_222 nm_ /Θ_208 nm_ < 1) (Fig. 1b). An analysis of secondary structure based on main chain NMR chemical shifts indicated that the residue-specific helical propensity (p_Hel_) is larger than 0.9 across 9 contiguous residues for (P3-7)_2_ and larger than 0.95 across 16 residues for (P3-7)_3_. As it is not common for monomeric peptides to cooperatively fold into ɑ-helices in the absence of tertiary interactions, we verified their monomeric state by SEC-MALS and native mass spectrometry (MS) (Figs. S1d and S1e). In addition we removed the N-terminal PGAS motif, that can facilitate helix nucleation^11^, from these peptides (Fig. 1a) and observed that the *un*capped counterparts, *u*(P3-7)_2_ and *u*(P3-7)_3_, also have high helical propensity (Fig. S1f).

Finally we investigated the thermal stability of the helices by CD spectroscopy at temperatures up to 368 K (Fig. 1c). The CD spectra at 278 K were equivalent to those obtained upon cooling after thermal unfolding, indicating that the unfolded state of the peptide is soluble under our experimental conditions. We also characterized the structural properties of (P3-7)_3_ by NMR at physiological temperature, 310 K. A comparison of the 13C-detected 2D CACO spectra of (P3-7)_3_ at 278 and 310 K revealed only a small decrease in helical propensity (Fig. 1c, center and right) indicating that the peptide remains essentially fully folded at 310 K (p_Hel_ ≈ 0.90 over 12 contiguous residues). In conclusion, concatenating Gln_i+4_→Leu_i_ interactions allows obtaining stable ɑ-helices that remain folded under physiological conditions, are soluble upon thermal unfolding and only contain natural amino acids.

### Structure of a Gln-based single ɑ-helix

The high quality of the NMR spectra obtained for peptide (P3-7)_2_ allowed using this technique to study its structure at atomic resolution and further characterize the interactions stabilizing its conformation (Fig. S2a). First we measured ^15^N relaxation at 278 K (R_1_, R_2_, heteronuclear ^15^N{^1^H} NOE) at two magnetic field strengths (14.1 T and 18.8 T) for the main chain amide (NH) groups of residues Gly4 to Lys24 and for the side chain amide (N_ε2_H_ε21_) groups of all four Gln residues (Gln11, Gln14, Gln17, Gln20) (Fig. 2a, S2b). We found that the main chain ^15^N R_2_/R_1_ ratios increase from the termini towards the center of the peptide until reaching plateau values (5.05 ± 0.25 at 18.8 T, 3.20 ± 0.15 at 14.1 T) between residues Leu10 and Gln17 (Fig. 2a). A similar trend is traced out by the ^15^N{^1^H} NOE, that reaches upper plateau values between 0.65 and 0.83 over a larger central segment, from Leu7 to Ala18. The main chain amide ^15^N relaxation data thus localizes the region of highest structural rigidity on a 10^-1^ to 10^1^ ns timescale as the one defined by the pairs of Leu and Gln residues involved in concatenated Gln_i+4_→Leu_i_ interactions *i.e.* from Leu10 to Gln17. By contrast, the Gln20→Leu16 interaction, involving the last Gln residue, appears to be weaker.

**Fig. 2.**
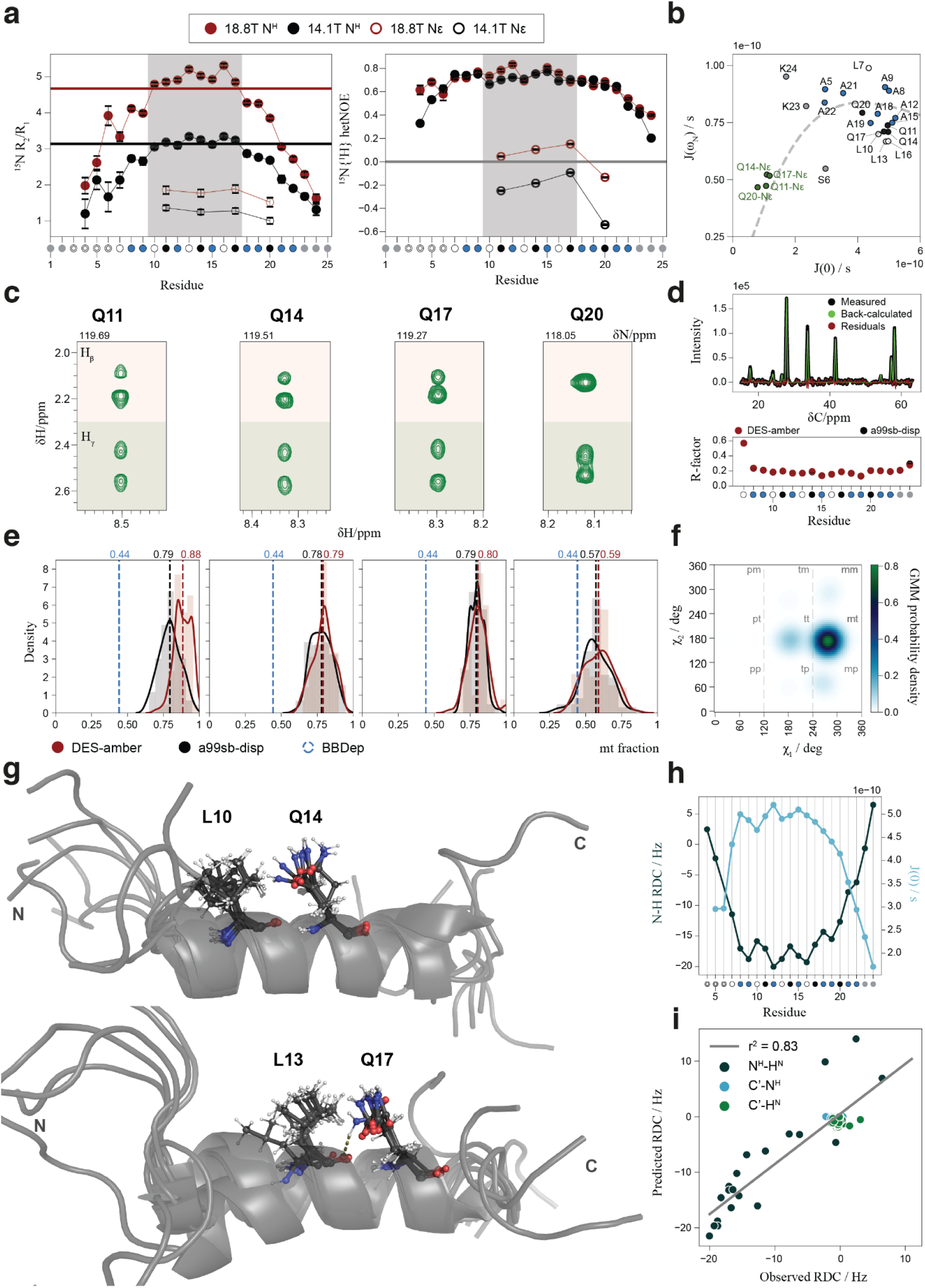
Structure of a Gln-based single ɑ-helix. **a** ^15^N NMR relaxation data for peptide (P3-7)_2_ main chain (N^H^) and Gln side chain (Nε) amide groups, measured at 14.1 T (black) and 18.8 T (dark red) at 278 K in 10% D_2_O (main chain) and 50% D_2_O (side chain), where residues are labeled as in Fig. 1. Left: ^15^N R_2_/R_1_ ratios. Solid horizontal lines represent the theoretical R_2_/R_1_ ratio of a rigid body in isotropic motion with τ_c_ = 4.5 ns, whose field dependency agrees well with the experimental data. Right: ^15^N{^1^H} heteronuclear NOE, where the shade corresponds to the region with the highest degree of structuration. **b.** Spectral densities, J(ω), derived by reduced spectral density mapping of all ^15^N relaxation data: the dashed line corresponds to the behavior expected for a polypeptide. **c** Strips from a 3D ^15^N-edited TOCSY-HSQC spectrum showing the H_β_ and H_γ_ resonances of all four Gln residues in peptide (P3-7)_2_. **d** Residue-specific CoMAND fitting of NOE signal intensities from the 3D CNH-NOESY (i.e. [H]C,NH HSQC-NOESY-HSQC) spectrum of peptide (P3-7)_2_. Top: comparison of measured and back-calculated (using CoMAND and the a99sb-disp trajectory frame pool) 1D ^13^C traces for the Gln14 main chain N. Bottom: Average R-factors obtained from 100 CoMAND iterations using either DES-amber or a99sb-disp trajectory frame pools. **e** Density plots of the fraction of the *mt* rotamer found in the 100 fitting iterations for all four Gln residues. The fraction of helical glutamines in the *mt* configuration in the BBDep dataset^26^ is shown in blue and marked with a blue dashed line, whereas black and dark red values and the corresponding dashed lines indicate the mean *mt* content for the a99sb-disp and DES-amber distributions, respectively. **f** Probability density for Gln14 χ_1_ and χ_2_ torsion angles derived by CoMAND using the Gaussian mixture model (GMM). **g** Structural ensemble of (P3-7)_2_, with main chain conformations selected by CoMAND from both trajectory pools and side chain conformations generated by GMM sampling. Top: one of the 20 global ensemble calculations aligned for the Leu 10 Gln14 pair. Bottom: same iteration aligned for the Leu13 Gln17 pair. **h** Residual dipolar couplings (RDCs) for the main chain ^1^H-^15^N moieties (dark blue, left axis) and relaxation-derived main chain spectral density at zero frequency, J(0), both showing the dipolar wave modulation characteristic of ɑ-helices. **i** Correlation between experimental and back-calculated RDCs, computed as the average of the ensemble produced by pooling the ensembles obtained in 20 independent iterations.

The different properties of the first three Gln residues (11, 14 and 17) relative to the last Gln residue (20) are also evident in the side chain amide ^15^N_ε2_ relaxation data: while positive (at 18.8 T) or close to zero (at 14.1 T) ^15^N_ε2_{^1^H_ε21_} NOE values suggest significantly less side chain mobility for the first three Gln residues, Gln20 shows clearly negative values in line with higher side chain mobility (Fig. 2a), also reflected in the relaxation-derived spectral density map (Fig. 2b). These results agree with the notion that the first three Gln residues form stronger Gln_i+4_→Leu_i_ interactions than Gln20. The high NMR signal dispersion in both the ^13^C and ^15^N dimensions of the spectra of peptide (P3-7)_2_ also allowed studying the Gln side chains: the first three Gln residues show fully resolved H_β_ and H_γ_ signals in the ^15^N-edited HSQC-TOCSY spectrum, whereas the H_β_ signals overlap for Gln 20 (Fig. 2c). This indicates that the conformations of the side chains of the former are better defined that those of the latter, even more than in polyQ helices^11,13^, in agreement with the relaxation data.

To improve the description of the Gln side chains, we used CoMAND^27^ to infer rotamer populations from a diagonal-free 3D CNH-NOESY spectrum reporting on distances between protons bound to ^15^N and ^13^C. Briefly, CoMAND selects subsets of conformers from a pool, here obtained by molecular dynamics, that reproduce the NOESY spectra (Fig. 2d). The distributions were enriched in the *mt* Gln rotamer (χ_1_ = -60° and χ_2_ =180°) that is required for the Gln_i+4_→Leu_i_ interaction: the Gln residues involved in strong interactions (11, 14 and 17) have a high population (0.80), whereas Gln 20 has a lower population (0.58). Both values are higher than that obtained for Gln residues in ɑ-helices of structures deposited in the PDB^26^, 0.44 (Fig. 2e); these results were robust to changes in the force field used to generate the pool^28,29^.

To generate a conformational ensemble, we used the residue-specific CoMAND ensembles to train a Gaussian mixture model (GMM) by inferring χ_1_ and χ_2_ probability densities for each residue (Figs. 2f and S2c), modified the side chain conformations accordingly and, through R-factor minimization, obtained the set of representative conformers shown in Figure 2g. To validate it we measured three sets of residual dipolar couplings (RDCs) under steric alignment. The main chain ^1^D_H,N_ values show a dipolar wave pattern typical of helices that matches well the period of the zero frequency spectral density, J(0) (Fig. 2h). Although the RDCs were not used as restraints they correlate well with the ensemble-averaged values (Q = 0.37, Fig. 2i), confirming that the ensemble is an accurate representation of peptide (P3-7)_2_ and that the design rules that we put forward lead to single ɑ-helices.

### Gln side chain to main chain hydrogen bonds can be accepted by residues other than Leu

The stability of the Gln-based ɑ-helices stems from concatenated Gln side chain to main chain hydrogen bonds accepted by Leu residues. This design decision was based on the fact that the polyQ tract found in AR, which is the most helical studied so far, is flanked by four Leu residues^11^. We sought to determine how other residues perform as acceptors to increase the versatility of the design rules and better understand the factors determining the strength of the interaction. For this we used a host-guest approach in which we determined the secondary structure of L_3_XQ_16_ peptides (Table S1) by NMR. These peptides were obtained by mutating the fourth Leu residue of peptide L_4_Q_16_, excised from AR, to 13 representative amino acids (Fig. 3a).

**Fig. 3.**
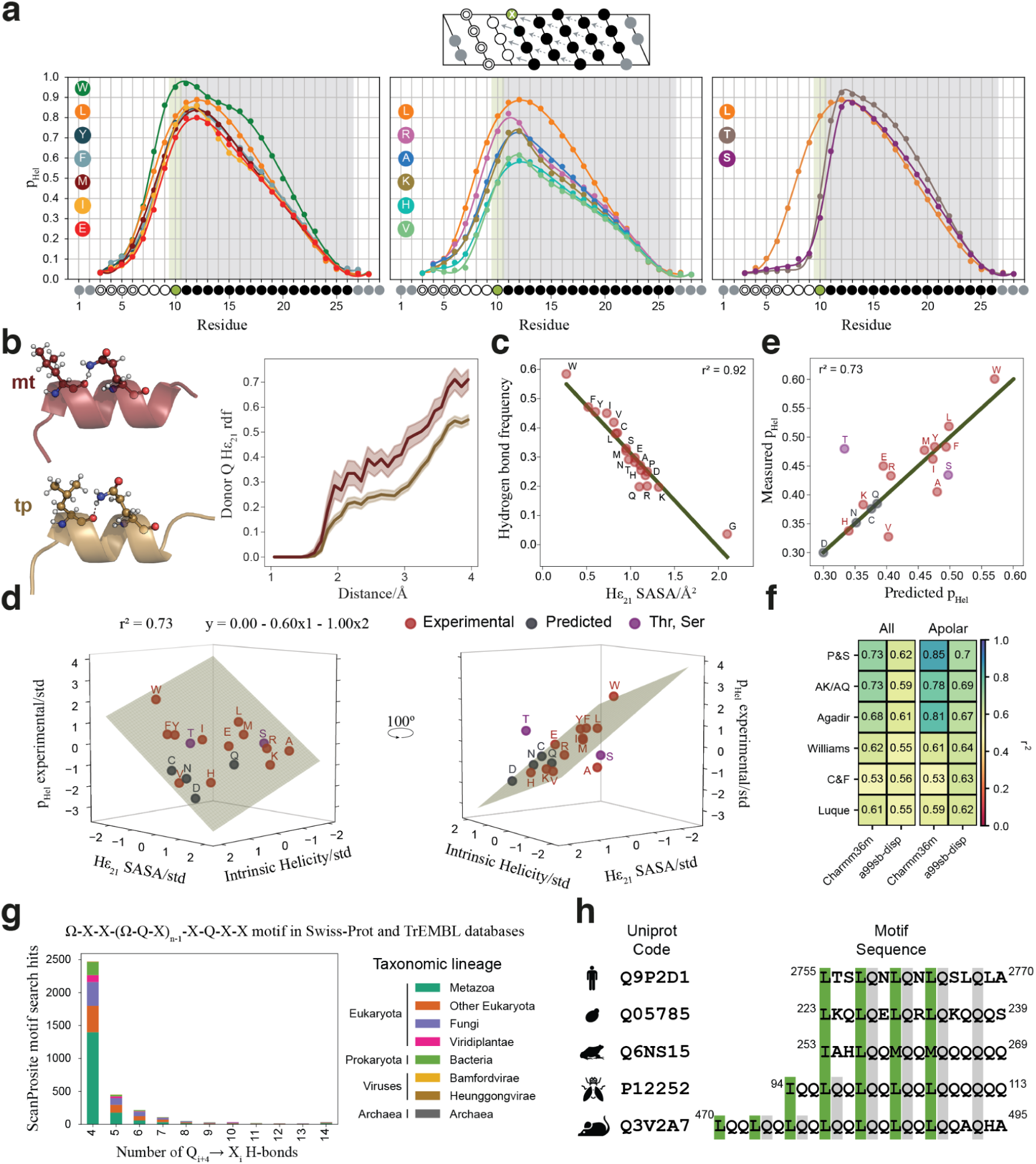
Gln side chain to main chain hydrogen bonds can be accepted by different residues. **a** Top: Representation, as helical projection, of the L_3_XQ_16_ variants studied in this work with the color code described in Fig. 1. Bottom: residue-specific helical propensity of the L_3_XQ_16_ variants derived from the backbone N, H^N^, C_α_ and C’ chemical shifts. The type of residue X (in position 10) is indicated by the colored circles. Left: helical profile of the 6 most helical single variants (X = W, Y, F, M, I, E) compared to that of L_4_Q_16_ . Center: helical profile of the least helical single variants (X = R, A, K, H, V) compared to that of L_4_Q_16_. Right: Helical profile of the outlier variants (X = T, S) compared to that of L_4_Q_16_ (spectra in Fig. S3b). **b** Effect of the rotameric state of the acceptor on the interaction of atom H_ε21_ with H_2_O. Left: two frames of the Charmm36m 1 μs MD trajectory obtained for L_4_Q_8_ showing Gln4 involved in a bifurcated hydrogen bond with Leu4 in either the *mt* (top) or *tp* (bottom) rotamer of the Leu residue. Right: radial distribution function for Gln4 H_ε21_ in the frames where Leu4 populates the *mt* (red) or *tp* (gold) rotamer. **c** The frequency of the side chain to main chain hydrogen bond is strongly correlated with the solvent accessibility surface area (SASA) of H_ε21_, which depends on type of residue X. **d** Multiple regression correlating intrinsic helicity (x1) (Pace and Scholtz scale^30^) and SASA (x2) with the average helicity (y): the data were standardized to estimate the relative weight of each variable in defining the model. The experimental data points used for regression are shown in dark red, the outliers in purple, and the model-predicted data points in teal. **e** Measured *versus* predicted average L_3_XQ_16_ helicity. Color code as in **d**. **f** Squared r correlation scores (r^2^) for the multiple regression shown in **e**, using different reported scales for intrinsic amino acid helicity (y axis) and H_ε21_ SASA values derived from two sets of 1 μs MD trajectories independently generated with different force fields. The results are shown for all experimentally measured variants (left) and those with an apolar residue (L, I, V, F, Y, M, W, A) in position X (right). **g** Number of ScanProsite-identified protein sequences in UniprotKB (including the Swiss-Prot and TrEMBL databases) containing the (P3-7)_n_ motif with an increasing number of Q_i+4_ → Ω_i_ pairs (Ω-X-X-(Ω-Q-X)_n-1_-X-Q-X-X motif search), where Ω represents any good side chain to main chain hydrogen bond acceptor W, L, Y, F, I and M, whereas X represents any amino acid. Counts are grouped by taxonomic lineage as obtained from UniprotKB. **h** Representative natural sequences that fulfill the design rules identified in proteins with UniprotKB annotation score = 5 and for which there is evidence at the protein level, belonging to a variety of organisms (from top to bottom: *Homo sapiens, Saccharomyces cerevisiae, Xenopus laevis, Drosophila melanogaster, Mus musculus*).

We measured the residue-specific helical propensities of the peptides by NMR by combining standard ^1^H^N^-detected triple resonance with ^13^C-detected CACO and CON 2D NMR experiments: the high resolution in the CO dimension of the latter allowed the unambiguous assignment of all Gln residues, even in the variants with lowest signal dispersion (Fig. S3b). Except L_3_TQ_16_ and L_3_SQ_16_ all variants show a helicity profile approximately proportional to that of L_4_Q_16_, (Fig. 3a): L_3_TQ_16_ and L_3_SQ_16_ instead show a different profile, likely because in these cases the substitution shifts the site of helix nucleation by introducing an S/T N-capping motif^31^. In these, Ser/Thr accept two concomitant hydrogen bonds donated by main chain amides of residues C-terminal to them: one by the main chain O and another one by the side chain O of the hydroxyl group.

To explain the range of helicities obtained we hypothesized that it is due to two main factors: the intrinsic helical propensity of residue X^30^ and, based on our previous work^11^, the ability of its side chain to shield the hydrogen bond. While the former has been extensively measured^23,30,32^, we quantified the latter by using molecular modeling, considering that the conformation of the residue accepting the hydrogen bond (X_i_) can have an effect on the interaction of the Gln_i+4_ H_ε21_ donor with competing water molecules (Fig. 3b and S4a). For this we computed 1 μs trajectories in different force fields^29,33^ for all 20 possible variants in which we constrained the secondary structure of the Leu2-Gln5 segment and increased the population of the Gln_i+4_→Leu_i_ interaction with a soft restraint to facilitate sampling the relevant region of conformational space: we obtained, as expected, that the higher the frequency of the hydrogen bond, the lower the solvent accessible surface area (SASA) of the H_ε21_ atom of Gln 4, with high correlation (Fig. 3c).

Next, we quantified to what extent intrinsic helical propensity (x1) and Gln_i+4_ H_ε21_ SASA (x2) explain the experimental helical propensities (Fig. 3d) by multiple linear regression. Indeed we obtained that these two independent variables explain 73% of the variability (Fig. 3d). Remarkably, the Gln 4 H_ε21_ SASA value (x2) is the most important factor in the correlation, as its weight in the fitted equation is 40% higher than that of intrinsic helicity (x1); the model allows the prediction of the average helicities of the L_3_XQ_16_ variants not included in our experimental dataset (Fig. 3e). The correlation improves (from r^2^ = 0.73 to 0.85) when only the subset of residue types with apolar side chains is considered, suggesting that additional factors might play a role when charged or polar side chains are present (Fig. 3f), and the results are robust to changes in MD force field^29,33^ or intrinsic helical propensity scales (Fig. 3f). These data confirm that Leu is one of the best helicity-promoting acceptors, but that other residues such as Phe, Tyr, Ile or Met are similarly good, and that Trp is an outstanding acceptor despite its relatively low intrinsic helical propensity. Thus, residues other than Leu can be introduced as acceptors of Gln side chain to main chain interactions, increasing the versatility of our design rules.

These results prompted us to investigate the presence of (P3-7)_n_-like motifs in nature. To do that, we searched UniprotKB^34^, including the Swiss-Prot and TrEMBL databases, by using the motif search tool in ScanProsite^35^, which we queried with the Ω-X-X-(Ω-Q-X)_n-1_-X-Q-X-X motif with Ω denoting good acceptors of Gln side chain to main chain hydrogen bonds (namely W, L, F, Y, I, M). We found that 3451 proteins contain sequences matching the motif, belonging to organisms across the kingdoms of life with representatives of a wide variety of taxonomic lineages including archaea, bacteria, viruses and a full range of eukaryotes, from unicellular organisms to metazoa including humans (Fig. 3g). There is experimental evidence for the existence of 94 of these proteins (UniprotKB annotation score > 3), mostly belonging to extensively characterized metazoa (Fig. S4b). Fig. 3h shows an alignment of some example sequences along with their UniprotKB accession code and the organism they belong to. We also found structural models for 50 of these sequences in the AlphaFold Database (AFDB)^18^: in 52% of the cases, a DSSP analysis^36^ of the AlphaFold model shows that the (P3-7)_n_-like motif is helical, increasing to 76.5% when only motifs devoid of helix breaking residue types (P, G) in the central part of the sequence are considered (Fig. S4c). In summary these findings suggest that (P3-7)_n_-like motifs may represent a new class of naturally occurring single α-helices (SAHs).

### Glu residues as pH-sensitive conformational switches

Gln to Glu substitutions in polyQ helices decrease helical character due to the inability of the Glu side chain to donate hydrogen bonds at physiological pH, where the carboxylate group is deprotonated^11^ (Fig. 4a). Re-protonation by decreasing the pH can restore the interaction, providing us with an opportunity to introduce a pH-sensitive conformational switch in our design rules. To explore this possibility we first sought to establish whether the loss of helicity upon substitution and its restoration upon pH decrease is strictly local or whether its effects instead can propagate to other parts of the sequence due to cooperativity.

**Fig. 4.**
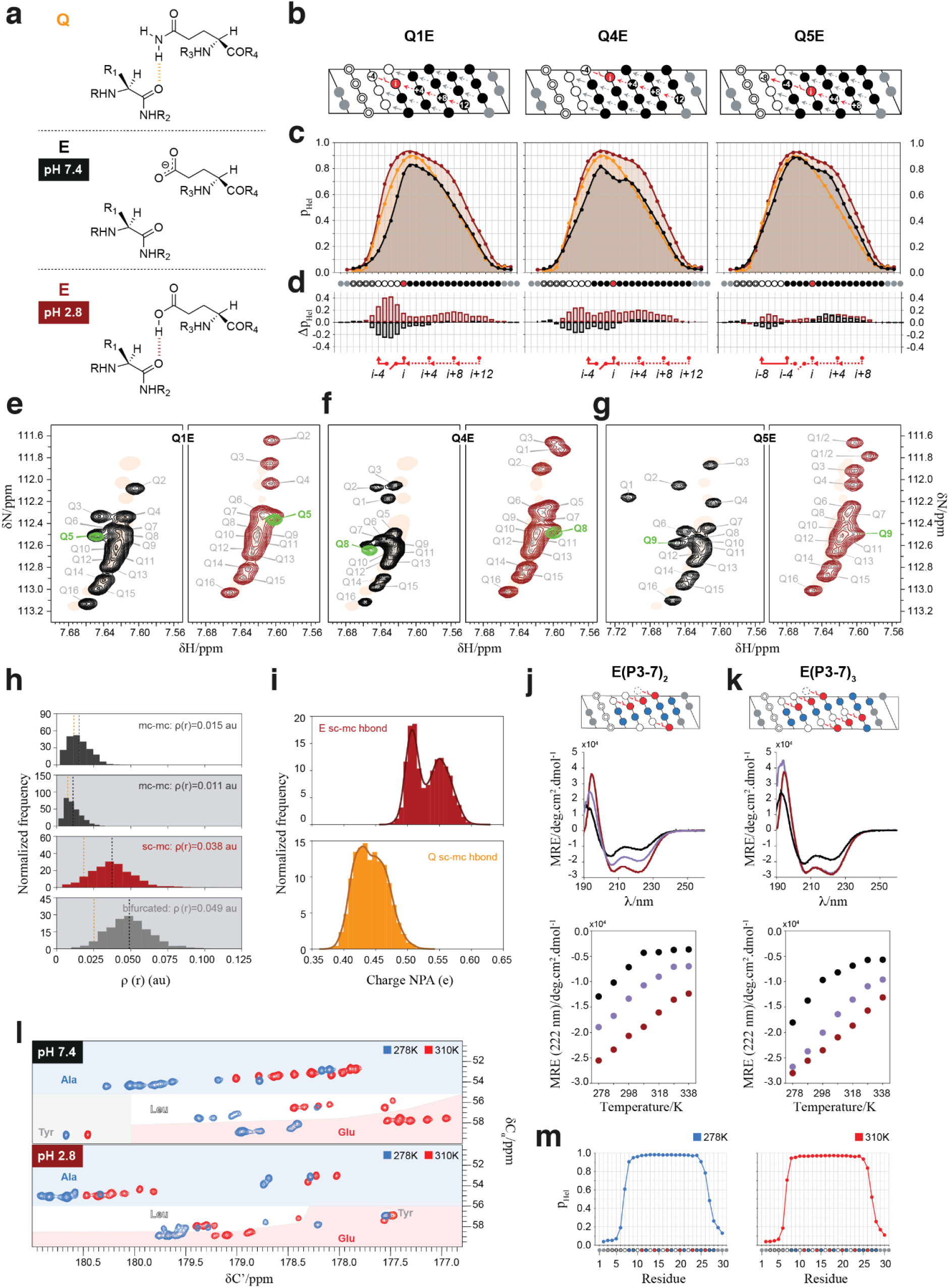
Introduction of a pH-sensitive conformational switch. **a** Schematic representation of how Gln (top) and Glu at pH 2.8 (bottom), but not Glu at pH 7.4 (center) side chains can donate a hydrogen to a main chain CO. **b** Representation as helical projection of the QxE variants of L_4_Q_16_ with pH-sensitive Glu_i+4_→X_i_ interactions shown as red switches and the residue substituted by E as a red circle; the color code is described in Fig. 1. **c** Residue-specific helical propensities at pH 7.4 (black) and 2.8 (dark red) compared to those of L_4_Q_16_ at pH 7.4 (orange). **d** Residue-specific differences in helical propensity due to substitution of Gln by Glu at pH 7.4 (black) and pH 2.8 (dark red). **e to g** Gln side chain N_ε2_-H_ε21_ regions of the 2D ^1^H-^15^N HSQC NMR spectrum of QxE variants at pH 7.4 (left panel, black) and 2.8 (right panel, dark red) with, for reference, the spectrum of L_4_Q_16_ as an orange shade. The spectrum of variants Q1E and Q4E with ^15^N labeling of only the Gln in position *i+4* to the mutated residue is superimposed in green as a verification of the assignment. **h** QM/MM-derived hydrogen bond electron densities. For reference, mean values for the Gln_i+4_→Leu_i_ interaction^11^ are shown as an orange dashed line. First and second panels: normalized histograms showing the distribution of the electron density *ρ(r)* of the main chain to main chain Glu_i+4_➝Leu_i_ hydrogen bond in the absence (white background) and in the presence (gray background) of the side chain to main chain Glu_i+4_➝Leu_i_ hydrogen bond. Third panel: distribution of the electron densities for the side chain to main chain Glu_i+4_➝Leu_i_ hydrogen bond. Fourth panel: distribution of the electron densities for the bifurcated hydrogen bond, resulting from the addition of the second and third panels. **i** Natural population analysis (NPA) charges on the hydrogen atom that can participate in the side chain to main chain hydrogen bond involving Glu (top) and Gln (down), showing that its charge depends on whether it participates in that hydrogen bond (lower values) or not. The clear peak separation in the bimodal distribution observed for Glu reports a higher electron polarization upon side chain to main chain hydrogen bonding when compared to Gln. **j to k** pH-dependent helicity and thermal stability for peptide E(P3-7)_n_, where all Gln residues in (P3-7)_n_ were substituted by Glu. Top: Helical projection for the peptide with an indication of all Gln to Glu substitutions (red). Center: CD spectra (at 278 K) of peptides E(P3-7)_n_ at pH 7.4 (black) or pH 2.8 (dark red) and of (P3-7)_n_ at pH 7.4 (purple). Bottom: thermal denaturation of peptide E(P3-7)_n_ (same pH and color code as above) monitored by measuring the mean residue ellipticity at 222 nm. **l** 2D CACO NMR spectra of [U-^13^C] labeled peptide E(P3-7)_3_ acquired at pH 7.4 (above), pH 2.8 (below), 278 K (blue), or 310 K (red). **k** Residue-specific helical propensities for peptide E(P3-7)_3_ at pH 2.8 and 278 K (left) or 310 K (right).

For this, we compared the residue-specific helicity of polyQ L_4_Q_16_ variants with Gln to Glu substitutions at positions 1 (Q1E), 4 (Q4E), and 5 (Q5E) in the polyQ tract (Fig. 4b and S4a). In Q1E the positions experiencing the strongest decrease in helicity are Leu 2 to Leu 4, which are tethered by the first bifurcated Gln_i+4_→Leu_i_ hydrogen bond in L_4_Q_16_, as expected, but there is also a small decrease in helicity for Gln residues in positions 5 to 8 (Fig. 4d, left, black). Similar effects were observed in Q4E and the loss of helical character in Q5E was much smaller, likely because the interaction broken upon substitution, accepted by Gln1, is weak even at physiological pH. This indicates that breaking the first Gln_i+4_→Leu_i_ interaction also weakens the interaction to which it is concatenated in the polyQ helix, and vice-versa, again in line with the notion that these interactions form cooperatively (Fig. 4c and d, black).

Experiments at pH 2.8, where the carboxylate group of the Glu side chain is protonated, showed that helicity was even higher than that of peptide L_4_Q_16_ at physiological pH (Fig. 4c and d, red); in addition the dispersion of Gln side chain N_ε_ and H_ε21_ resonances in the 2D ^1^H,^15^N HSQC spectrum, that is characteristic of polyQ helices, was restored (Fig 4e-g). To investigate the physical basis of this, we simulated this interaction by using a hybrid QM/MM approach (Fig. S5b and c). We found that the establishment of the side chain to main chain hydrogen bond weakened the main chain to main chain hydrogen bond: the associated average electron density decreased from 0.015 a.u. to 0.011 a.u. (Fig. 4h) and the main chain Glu_i+4_ (H) - Leu_i_ (O) interatomic distance increased by 0.15 Å (Fig. S5d). Consistent with the experimental stronger hydrogen bond donor character of protonated Glu, relative to that of Gln, the average electron density of the Glu_i+4_ ➝ Leu_i_ side chain to main chain hydrogen bond (0.038 a.u.) was higher than that involving Gln side chains (0.017 a.u.) and the electronic polarization of the H_ε2_ donor was stronger for Glu relative to Gln (Fig. 4i).

Finally, to test whether switchable side chain to main chain hydrogen bonds can be integrated in our design rules, we studied variants of the (P3-7)_2_ and (P3-7)_3_ peptides where all Gln residues were substituted by Glu, namely E(P3-7)_2_ and E(P3-7)_3_ (Figs. 4j and 4k, top). As expected, these were less helical than (P3-7)_2_ and (P3-7)_3_ at physiological pH, but more at pH 2.8 (Figs. 4j and 4k, center). Both the switchable nature and the increased strength of the side chain to main chain hydrogen bonds involving Gln is apparent in the enhanced thermal stability of the E(P3-7)_n_ variants (Figs. 4j and 4k, bottom): while at physiological pH peptide E(P3-7)_3_ loses its helical character at a lower temperature than (P3-7)_3_, at pH 2.8 it remains helical even at the highest tested temperature, 340 K. Indeed ^13^C-detected 2D CACO NMR spectra confirmed that, at pH 2.8, E(P3-7)_3_ was highly helical at 310 K, the physiological temperature (Figs. 3l and m).

### Gln_i+4_→X_i_ interactions can be combined with electrostatic interactions between side chains

The natural single ɑ-helices studied until now are stabilized by numerous electrostatic interactions between side chains of opposite charge at relative positions *i,i+3 or 4*^37^. We sought to investigate whether they can be combined with Gln_i+4_→X_i_ interactions to stabilize ɑ-helices. For this we studied the polyQ tract of the TATA-box binding protein (TBP), that has a primary structure that suggests the presence of an electrostatic interaction, between either of two Glu residues immediately flanking the tract at the N-terminus (Glu9 and Glu10) and an Arg interrupting it (Arg13) (Fig. 5a). This interaction can occur concomitantly with two strong bifurcated hydrogen bonds accepted by Ile7 and Leu8, at position *i-4* relative to the first two Gln residues of the tract. As observed for the polyQ tracts in AR ^11^ and huntingtin^38^ the CD spectrum of a peptide spanning a tract of size 16 and its N-terminal flanking region, TBP-Q_16_, showed it is is strongly helical and that its expansion to 25 Gln residues increases its helicity (Fig. 5a,b and S6).

**Fig. 5.**
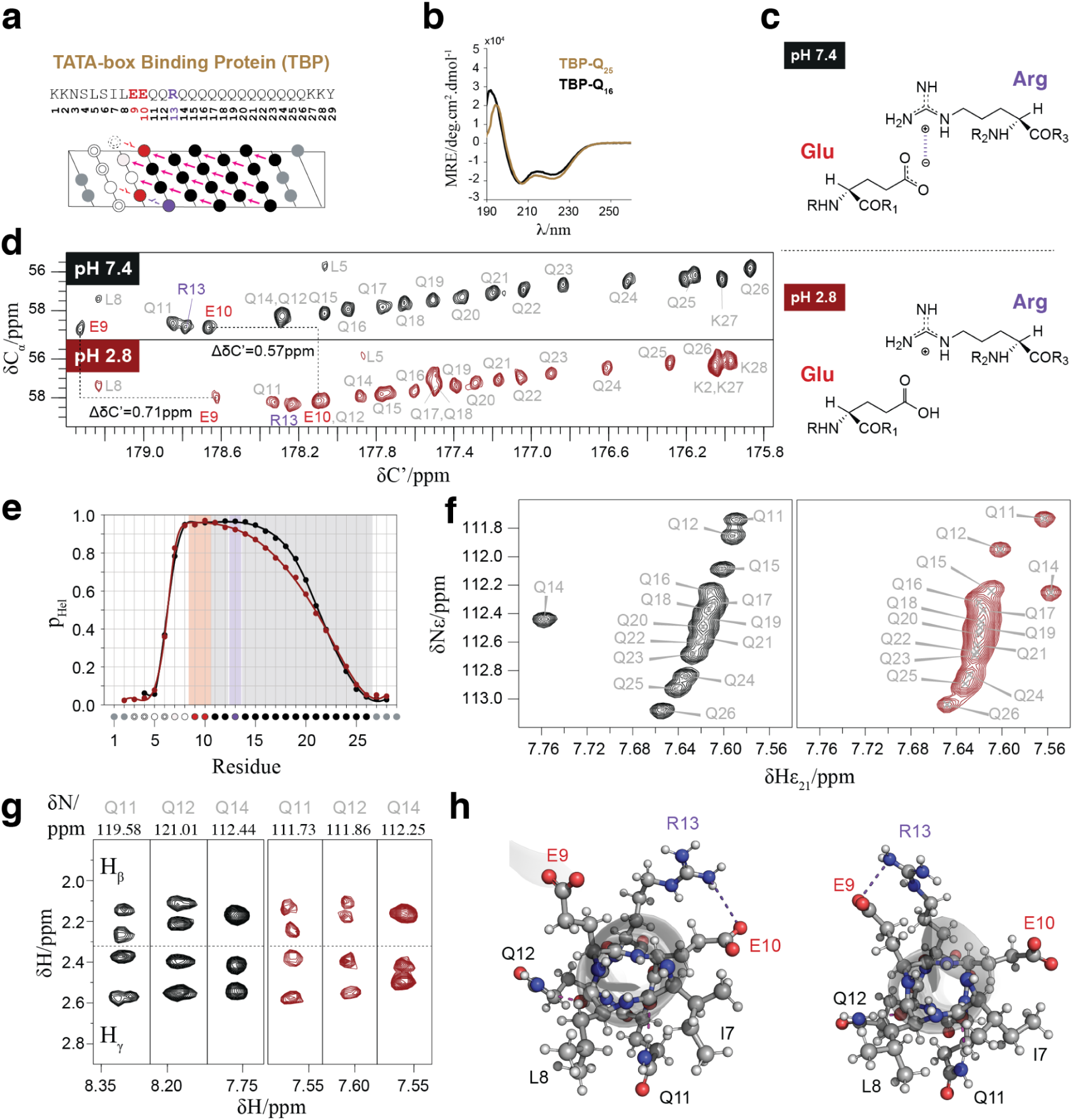
The polyQ tract of TBP forms a helix stabilized by Gln side chain to main chain hydrogen bonds and an electrostatic interaction . **a** Sequence, numbering and helical projection of peptide TBP-Q_16_. The electrostatic interaction is color-coded in purple. **b** CD spectra of peptides TBP-Q_16_ (black) and TBP-Q_25_ (gold). **c** Scheme of the electrostatic interaction between Arg and Glu at physiological pH and its absence at acidic pH. **d** ^13^C-detected CACO spectra of the TBP polyQ tract at pH 7.4 (top, black) and 2.8 (dark red, down). **e** Residue-specific helicity of TBP at pH 7.4 (black) and 2.8 (dark red). Shades color-coded as in **a** are shown to guide the eye. **f** Region of the ^1^H-^15^N HSQC spectra of TBP at pH 7.4 (left, black) and 2.8 (right, dark red) showing the N_ε2_-H_ε21_ correlations. **g** Strips from the 3D H(CC)(CO)NH spectra of TBP at pH 7.4 (left, black) and 2.8 (right, dark red) displaying the side chain aliphatic ^1^H resonances of the first three Q residues in the polyQ tract. Strips were chosen from either N^H^ or N_ε2_ for clarity. **h** Frames from the MD trajectory obtained for TBP. An orthogonal view of the helix is shown. In the left panel, a frame is shown where Arg13 (*i*) establishes an electrostatic interaction with E10 (*i-3*), represented by the purple dashed line. Simultaneously, both Gln11 and Gln12 establish bifurcated hydrogen bonds with Ile7 and Leu8, respectively (pink dashed lines). Right: a frame where Arg13 (*i*) establishes an electrostatic interaction with Glu9 (*i-4*), while the two bifurcated hydrogen bonds previously described are also present.

We then used NMR to characterize TBP-Q_16_ at residue resolution (Fig. 5d,e). At physiological pH, in agreement with the CD data, the peptide forms a fully folded helix between residues Glu9 and Gln14 and its helicity decreases progressively towards the C-terminus; at acidic pH, instead (Fig. 5c), at which Glu side chains are protonated, the helical propensity starts decreasing at position 12. In addition, both the spectral signature of concatenated Gln_i+4_→X_i_ interactions (Fig. 5f) and the rotamer selection associated with these interactions (Fig. 5g) are diminished for Gln14 (H_γ_) and, to some extent, Gln12 (H_β_). These results are in agreement with the formation of a helix-stabilizing electrostatic interaction between Glu9 (or Glu10) and Arg13 that is lost upon protonation at low pH, confirming that electrostatic interactions can be combined with Gln side chain to main chain interactions. To further confirm that these interactions can co-exist we simulated a WTE-enhanced MD trajectory of TBP-Q_16_^29,39^: Fig. 5h shows two frames of the trajectory in which both the Gln11➝Ile7 and the Gln12➝Leu8 bifurcated hydrogen bonds occur simultaneously with a salt bridge involving Arg13 and either Glu10 (left) or Glu9 (right).

### Design of Gln-based single α-helices that bind a globular target

Gln-based ɑ-helical peptides can be modified to bind specific globular targets: the Ala residues in the (P3-7)_n_ scaffolds (Fig. 1) can indeed be modified at will because they are not involved in the interactions that stabilize the helical structure. To prove this concept we modified the sequence of peptide (P3-7)_3_ to interact with the C-terminal domain of RAP74 (RAP74-CTD), a small globular domain that binds to intrinsically disordered motifs that fold upon binding^40–42^. We blended the sequence of (P3-7)_3_ with that of two different motifs (centFCP1 and cterFCP1) derived from FCP1 that interact with this globular protein independently^42^. This led to peptides δ and δ_ctrl_: δ was designed to bind to RAP74-CTD whereas δ_ctrl_ is a control sequence equivalent to δ where we replaced Leu by Ala that, despite having high helical propensity, are bad acceptors of side chain to main interactions, thus destabilizing helicity (Fig. 6a and S7).

**Fig. 6.**
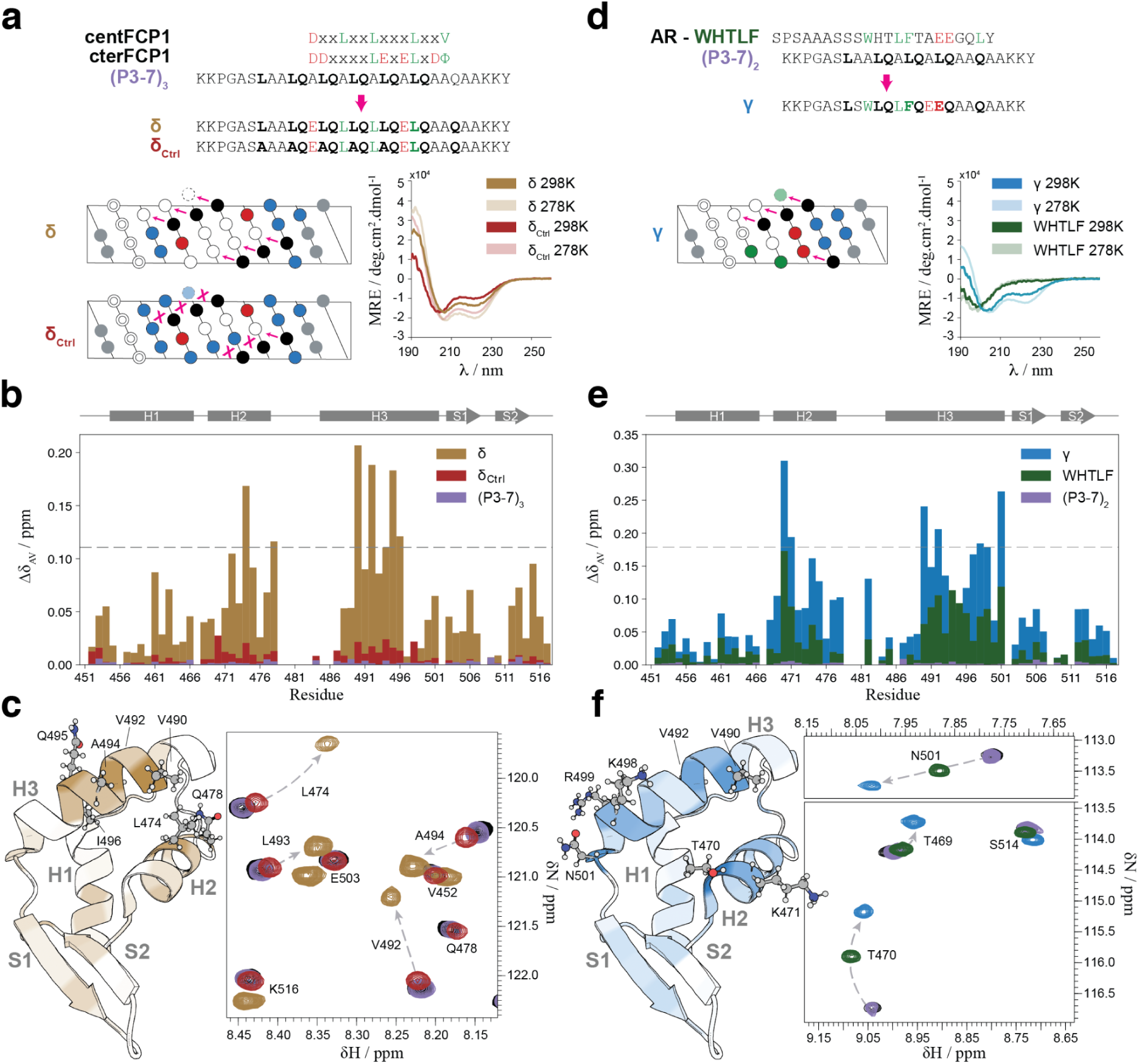
Design of Gln-based single helical peptides that interact with RAP74-CTD. **a** Top: design of peptide δ, blending the consensus RAP74-CTD binding motifs of FCP1 with peptide (P3-7)_3_. Peptide δ_ctrl_, in which Leu residues in δ are substituted by Ala, is also shown. Bottom, left: projections of the fully helical structures of peptides δ and δ_ctrl_. Bottom, right: CD spectra of peptides δ (gold) and δ_ctrl_ (dark red) at 298 K (saturated) and 278 K (pastel). **b** Averaged chemical shift perturbations observed in the ^1^H-^15^N resonances of the RAP74-CTD upon the addition of 15 molar equivalents of the (P3-7)_3_ (purple), the δ (gold) and the δ_ctrl_ (dark red) peptides. The dashed line sets the threshold for the top 10% peaks with most intense averaged CSPs. **c** Left: structure of the RAP74-CTD domain of TFIIF (PDB entry 1NHA) colored in a gradient representing the averaged CSPs observed upon the addition of 15 molar equivalents of the δ peptide (white, minimum CSP_av_; gold, maximum CSP_av_). Right: region of the overlaid ^1^H-^15^N BEST-TROSY spectra of RAP74-CTD in the absence (black) and in the presence of 15 molar equivalents of the (P3-7)_3_ (purple), the δ (gold) and the δ_ctrl_ (dark red) peptides. **d** Top: design of peptide γ blending the RAP74-CTD binding motif AR-WHTLF with peptide (P3-7)_2_. Bottom, left: projection of the fully helical structure of peptide γ. Bottom, right: CD spectra of peptide γ (blue) and AR-WHTLF (green) at 298 K (saturated) and 278 K (pastel). **e** as in **b** for (P3-7)_2_ (purple), peptide γ (blue) and peptide AR-WHTLF (green). **f** Left: as in **c**, left, for peptide γ. Right: as in **c**, right, for the (P3-7)_2_ (purple), the γ (blue) and the AR-WHTLF (green) peptides.

An analysis of the structural properties of peptides δ and δ_ctrl_ by CD showed, as expected, that the former is more helical than the latter, especially at room temperature (298 K), further confirming the important role of Leu residues for the stability of the helical fold (Fig. 6a). We then analyzed the chemical shift perturbations (CSPs) in the ^1^H,^15^N BEST-TROSY spectrum of RAP74-CTD induced by peptide binding. We obtained that δ induced perturbations in residues of the globular protein that define the binding site of FCP1 in this globular target^42^, confirming a similar binding mode. This was in contrast to the results obtained with both δ_ctrl_ and (P3-7)_3_, which in both cases failed to interact (Fig. 6b and c). The result obtained with δ_ctrl_ indicates that the helical character of δ is key for its ability to interact with RAP74-CTD whereas that obtained with (P3-7)_3_ indicates that helical character does not suffice, and that the identity of the residues placed in the vacant positions is indeed key for binding, in agreement with our hypothesis.

To provide a second proof of concept we blended the sequence of (P3-7)_2_ with that of a motif found in the activation domain of AR that also interacts with RAP74-CTD, to yield peptide γ (Fig. 6d). As we previously showed, inhibiting this interaction with small molecules or peptides is a potential avenue to treat castration-resistant prostate cancer^40^. Linear peptides spanning the AR motif bind weakly to RAP74-CTD due, at least in part, to their low helical propensity, providing us with an additional opportunity to test the potential of our designs. In peptide γ two Gln_i+4_➝X_i_ hydrogen bond acceptor positions were modified to accommodate the binding motif following the rules learned previously (Fig. 3). In agreement with our hypothesis we obtained that peptide γ binds more strongly than either (P3-7)_2_ or the WHTLF peptide (Fig. 6e and f). In summary we have shown that the Gln-based helical peptides can be adapted to interact with specific globular proteins.

## Discussion

### A new tool to design fully helical peptides

Our results show that Gln side chain to main chain hydrogen bonds can be used to design linear peptides that fold into α-helices (Fig. 2) with properties that make them attractive for various applications: they are highly soluble, even upon thermal denaturation, are not stabilized by electrostatic interactions, unlike the Glu and Lys/Arg-rich single α-helices reported until now^43–45;^ and display some degree of folding cooperativity due to the concatenation of side chain to main chain hydrogen bonds explicit in our design rules (Fig. 1). Our data shed light on the potential basis of such cooperative effect, which occurs when the donor of a Gln_i+4_→Leu_i_ interaction and the acceptor of the next one share a peptide bond.

An important feature of our design rules is their versatility: the residue accepting the hydrogen bond donated by the Gln side chain can be any residue able to shield the interaction from competition with water, such as Trp, Leu, Phe, Tyr, Met and Ile (Fig. 3). Remarkably, we find natural sequences fulfilling our design in different kingdoms of life suggesting that they represent a new class of uncharged single α-helices (SAHs) that had remained undetected^43–45^. In addition, in suitable cases the design can be complemented by electrostatic interactions between side chains of opposite charge and Gln residues can be mutated to Glu to introduce a pH-responsive conformational switch that uses only natural amino acids^46^ and does not involve changes in oligomerization state^47^ (Figs. 4 and 5).

The key feature of our scaffold design is that it defines the identity of just a fraction of the peptide residues: the rest can be chosen or optimized for specific applications. To prove this concept, we designed two peptides to bind the globular target RAP74-CTD by using an approach analogous to previous motif-grafting attempts on folded scaffolds^48^ (Fig. 6). In this specific case, naively blending these sequence features with the sequence of the designed (P3-7)_n_ scaffold proved sufficient for successful targeting. The affinities that we have obtained in this initial exercise (Fig. S6d and e) can be sufficient for certain applications^49^ and we anticipate that it will be possible to greatly improve them by systematically searching the sequence space available using techniques for affinity maturation based on high-throughput mutational scans^50^, especially when taking advantage of the versatility of our design rules.

### A rationale for the structural heterogeneity of polyQ tracts

The link between polyQ tract expansion and disease onset in polyQ disorders has not been established^51^. These tracts are found in intrinsically disordered regions, and much work has been devoted to investigating whether expansion changes their structural properties. The results obtained have been inconclusive: single molecule Förster resonance energy transfer (FRET) and NMR measurements showed little influence of tract length on the conformation of the tract found in huntingtin^13,52^ whereas recent studies from some of us have instead shown that the helical propensity of the tract found in the androgen receptor (AR) increases upon expansion^11,12^, as does the tract found in TBP (Fig. 5b).

Our results help rationalize these observations by considering that polyQ tracts are in a polyQ helix-coil equilibrium. Its position can be influenced by the residues flanking the tract at its C-terminus^13,53^ but, for a given set of solution conditions and tract length, it is mainly determined by the four residues flanking the tract at its N-terminus (Fig. 3). When these are good acceptors of Gln side chain to main chain hydrogen bonds, as in AR (LLLL), the polyQ helix is favored. Instead, when only two are good acceptors as in huntingtin (LKSF), the coil is favored^13,19,52,54^. The results obtained for TBP also fit this working model: its flanking region (ILEE) contains two good acceptors (Ile, Leu) and the two other residues (Glu) can establish, at physiological pH, an electrostatic interaction with the Arg residue three or four positions towards the C-terminus, favoring the polyQ helix.

The polyQ helix-coil equilibrium is sensitive to solution conditions: the entropic cost of folding results in higher stability of polyQ helices at relatively low temperatures whereas high temperatures favor the coil. This contributes in part to explaining the discrepancy between the results obtained with huntingtin, carried out at room temperature^52^, and those obtained for the androgen receptor, carried out at 5°C^11^: indeed, CD studies of the structural properties of huntingtin showed an increase in helical propensity upon tract expansion at low temperature (−10°C)^38^. Finally our observation that pairs of concatenated side chain to main chain interactions form cooperatively contributes to explaining how expansion shifts the equilibrium to the polyQ helix state both in AR^11^ and in huntingtin^52^.

In summary our results suggest that polyQ tracts are in a helix-coil equilibrium that is governed by the N-terminal flanking region, by solution conditions and, due to cooperativity, by tract length. Given that interactions between low-populated helical conformations of huntingtin play a role in the early stages of its aggregation into amyloid fibrils^55,56^ we speculate that polyQ expansion leads to the onset of Huntington’s disease at least in part by stabilizing pre-nucleation oligomeric species that are on-pathway to aggregation.

### Concluding remarks

In summary we have shown how an appropriate concatenation of Gln side chain to main chain hydrogen bonds makes it possible, on the one hand, to design highly helical peptides that can be tailored to specific applications and, on the other hand, to rationalize the until now perplexing observations regarding the structural properties of polyQ tracts. We anticipate that the knowledge gained about this interaction will influence future developments in peptide design, particularly in the use of peptides as therapeutics, as well as contribute to better understanding the molecular basis of polyQ diseases.

## Methods

### Peptide sample preparation

Recombinant peptides with ^15^N or ^13^C,^15^N isotope enrichment were prepared as detailed elsewhere^57^. Synthetic genes coding for the peptide of interest and codon-optimized for expression in *Escherichia coli*, with an N-terminal His_6_-Sumo tag fusion and cloned into the pDEST-17 vector, were directly obtained from GeneArt (Thermo Fisher Scientific, Waltham, MA, USA). Genes coding for the L_4_Q_16_ variants in the L_3_X and QXE series (see Table S1) were obtained using the Q5 Site-Directed Mutagenesis Kit from New England Biolabs (Ipswich, MA, USA). Rosetta (DE3)pLysS competent cells (Novagen, Merck KGaA, Darmstadt, Germany) were used for expression in M9 medium containing ^15^NH_4_Cl and, where required, ^13^C-glucose (both from Cambridge Isotope Laboratories Inc., Tewksbury, MA, USA) as the sole nitrogen and carbon sources, respectively. All purification steps were performed at 277 K. Cell lysates in lysis buffer (20 mM Tris-HCl, 100 mM NaCl, 20 mM imidazole, pH 8.0) were purified by immobilized metal affinity chromatography (IMAC) using a HisTrap HP 5 mL column mounted on an Äkta Purifier System (GE Healthcare, Chicago, IL, USA). Fractions containing the His_6_-Sumo-peptide fusion in elution buffer (20 mM Tris-HCl, 100 mM NaCl, 500 mM imidazole, pH 8.0) were pooled and dialyzed overnight in lysis buffer while treated with ubiquitin-like specific protease Ulp1 (50 μg/mL). The His_6_-Sumo tag was removed with an additional IMAC step and the peptide-containing flow-through was dialyzed in ultrapure MilliQ water before lyophilization. Unlabeled synthetic peptides from solid-phase peptide synthesis were directly obtained from Genscript (Piscataway, NJ, USA) as lyophilized powder with >95% purity. Both recombinant and synthetic lyophilized peptides were dissolved in 6M guanidine thiocyanate (Merck KGaA, Darmstadt, Germany) and incubated overnight at 1250 rpm, 298 K, in a thermoblock. The sample was then injected in an Äkta Purifier System equipped with a Superdex Peptide 10/300 GL column (GE Healthcare, Chicago, IL, USA) equilibrated with ultrapure water, 0.1% trifluoroacetic acid. The fractions containing monomeric peptide were pooled and centrifuged at 386000 g for 3 h in an Optima TLX ultracentrifuge equipped with a TLA 120.1 rotor (Beckman Coulter, Atlanta, GA, USA). Orthophosphoric acid or sodium phosphate were added to a final concentration of 20 mM to adjust the pH to 2.8 or 7.4, respectively. The peptide concentration was determined by measuring the absorbance at 280 nm (extinction coefficients were calculated using the Protparam tool on the ExPASy website, https://web.expasy.org/protparam) before diluting the sample to the final experimental concentration.

### Protein sample preparation

Recombinant samples with ^15^N isotope enrichment of the C-terminal domain of subunit 1 of the general transcription regulator TFIIF (RAP74-CTD), spanning residues 450-517, were obtained as described previously^40^. A codon-optimized synthetic gene cloned in a pDONR221 vector was obtained from GeneArt (Thermo Fisher Scientific, Waltham, MA, USA) and subcloned into a pDEST-His_6_MBP vector obtained from Addgene. Rosetta (DE3)pLysS competent cells (Novagen, Merck KGaA, Darmstadt, Germany) were grown in MOPS medium with ^15^NH_4_Cl as the only nitrogen source (310 K, induction at OD_600_ = 0.7, 1 mM IPTG, harvesting 3 h after induction). Soluble fractions of cell lysates from sonication in lysis buffer (50 mM Tris-HCl, 1M NaCl, 10 mM imidazole, pH 8.0) were purified by IMAC and fractions containing the His_6_MBP-TEV-RAP74-CTD fusion were dialyzed at 277 K overnight against cleavage buffer (50 mM Tris-HCl, 200 mM NaCl, 0.5 mM EDTA, pH 8.0) in the presence of TEV protease (50 μg/mL). A second IMAC step was performed and the flow-through containing the RAP-CTD was loaded onto a HiTrap SP HP cation exchange column followed by a size exclusion chromatography step on a Superdex 75 GL 10/300 column equilibrated with NMR buffer (20 mM sodium phosphate, 0.1% trifluoroacetic acid, pH 7.4), both mounted in an Äkta Purifier System (GE Healthcare, Chicago, IL, USA). The sample was concentrated to 100 μM using an Amicon Ultra 15 mL centrifugal filter (Merck KGaA, Darmstadt, Germany).

### CD spectroscopy

Peptide samples for CD spectroscopy were diluted to a final concentration of 30 μM in a volume of 400 μL in either 20 mM phosphoric acid (pH 2.8) or sodium phosphate (pH 7.4) buffer. Spectra were obtained at 278 K (unless stated otherwise) in a Jasco 815 UV spectro-photopolarimeter with a 1 mm optical path cuvette using a data interval of 0.2 nm in the 190 - 260 nm range with a scanning speed of 50 nm min^-1^ and 20 accumulations. A blank spectrum acquired on the pertaining buffer under the same experimental conditions was subtracted from the sample spectrum. Thermal denaturation experiments were performed by acquiring a single accumulation spectrum with the same parameters at 10 K intervals, with a temperature ramp speed of 10 K min^-1^ and an equilibration time of 1 min.

### NMR spectroscopy

Peptide samples for NMR spectroscopy were diluted to a final concentration of 100 μM in a volume of 400 μL in either 20 mM phosphoric acid (pH 2.8) or sodium phosphate (pH 7.4) buffer with added 10% v/v D_2_O and 10 μM DSS for internal chemical shift referencing, then filled into Shigemi tubes (Shigemi Co. Ltd, Tokyo, Japan). All NMR experiments were recorded at 278 K (unless stated otherwise) on either a Bruker Avance III 600 MHz or a Bruker Avance NEO 800 MHz spectrometer, both equipped with TCI cryoprobes. Unlabeled synthetic peptides P1-5, P2-6, P3-7, P5-9 were characterized by two-dimensional homonuclear (TOCSY and NOESY) and heteronuclear (^1^H-^13^C HSQC) experiments. The TOCSY and NOESY mixing times were set to 70 and 200 ms, respectively. Water suppression was achieved by excitation sculpting^58^ using a 2 ms long Squa100.1000 selective pulse. For peptide backbone resonance assignment, using uniformly ^15^N,^13^C labeled peptides, the following series of 3D triple resonance BEST-TROSY^59^ experiments were acquired with 25% non-uniform sampling (NUS): HNCO, HN(CA)CO, HN(CO)CA, HNCA, and HN(CO)CACB. For some peptides we resolved assignment ambiguities by also acquiring a 3D (H)N(CA)NH spectrum. Furthermore, 2D ^13^C-detected CACO and CON experiments^60^ were measured. Data processing was carried out with qMDD^61^ for non-uniform sampled data and with NMRPipe^62^ for all uniformly collected experiments. Data analysis was performed with CcpNmr Analysis^63^. Determination of the residue-specific helical propensity (p_Hel_) from backbone chemical shifts H^N^, N^H^, C’ and C_α_ was performed using CheSPI^14^, which uses sequence and condition-corrected (temperature, pH) estimates for the reference random coil chemical shifts derived from POTENCI^15^. CheSPI was chosen over other algorithms as it intrinsically considers chemical shift changes derived from Glu side chain protonation, as the ∼0.6 ppm C’ chemical shift change reported before^64^ and observed in the TBP CACO spectra (Fig. 5d).

Side chain aliphatic ^1^H chemical shifts were obtained from 3D ^15^N edited TOCSY-HSQC (75 ms mixing time) and NOESY-HSQC (200 ms mixing time) spectra. Glutamine side chain resonances were assigned using complementary 3D H(CC)(CO)NH and (H)CC(CO)NH spectra recorded with 25% NUS and 14 ms C,C-TOCSY mixing. To further confirm the side chain N_ε_ assignments for Gln5 in peptide Q1E and Gln8 in peptide Q4E we recorded 2D ^1^H-^15^N HSQC spectra of synthetic unlabeled peptides with specific ^15^N labeling only in the positions of interest.

For the detailed analysis of sidechain rotamer distributions by the CoMAND approach, a 3D CNH-NOESY (i.e. 3D [H]C,NH HSQC-NOESY-HSQC) spectrum^65^ of [U-^13^C,^15^N] labeled (P3-7)_2_ was recorded at 800 MHz, 278 K, with 400 ms NOE mixing time and 64(^15^N) x 86(^13^C) x 2048(^1^H) complex data points corresponding to 15.2 x 104.8 x 6.3 Hz FID resolution. The final ^15^N-HSQC module employed sensitivity-enhanced coherence selection by gradients and band-selective flip-back of H^C^ polarization to enable its faster re-equilibration during a shorter total interscan delay (experimentally optimized as 0.6 s). For prior assignment of all aliphatic side chain ^1^H and ^13^C resonances and easy distinction between intra- and inter-residual NOE signals in the [H]C,NH HSQC-NOESY-HSQC spectrum, we furthermore recorded a set of 3D [H]CC[CA]NH TOCSY (11.3 and 22.6 ms FLOPSY8 mixing) and [H]CC[CO]NH TOCSY spectra (9 and 18 ms FLOPSY8 mixing).

To study glutamine side chain ^15^N relaxation, a ^15^N labeled (P3-7)_2_ sample was prepared in NMR buffer (pH 7.4). To avoid bias due to dipole-dipole cross-correlated relaxation within the N_ε2_H_2_ moieties and thus allow a direct comparison with the main chain NH data, we sampled only their 50% semi-protonated N_ε2_HD isotopomers in buffered 50% D_2_O and applied continuous deuterium decoupling during the ^15^N coherence evolution. Of note, the differential deuterium isotope shift of ^15^N_ε2_ in the N_ε2_H_ε21_D_ε22_ *vs* N_ε2_D_ε21_H_ε22_ species^66^ also allowed an unambiguous stereospecific signal assignment of the attached side chain carboxamide H_ε_ (Fig. S2b).To measure ^15^N R_1_ and R_2_ rates the conventional pulse sequences with sensitivity-enhanced coherence selection by gradients, water flip-back, and fully interleaved acquisition of relaxation delays were complemented with continuous deuterium decoupling during t_1_(^15^N) in order to suppress ^15^N(t_1_) line broadening from scalar relaxation (via ^1^J_15N,D_ coupling) for the glutamine sidechain NHD isotopomers of interest. In contrast, the pulse sequence for measuring the ^15^N{^1^H} heteronuclear NOE (likewise with sensitivity-enhanced coherence selection by gradients and continuous deuterium decoupling during t_1_(^15^N)) required further critical adaptations to suppress detrimental antiphase signal components (in F1(^15^N)) for the 50% glutamine NH_2_ isotopomers that impede a clean quantification of nearby NHD signals of interest. Thus, for the reference (non-saturated) spectrum, the first 90° ^1^H pulse in the sensitivity-enhanced reINEPT following t_1_(^15^N) had to be cycled (inverted) along with the receiver phase. For the H^N^ saturated spectrum, however, further antiphase contamination derives from some 4N_z_H^±^H’^±^ multiquantum coherence forming during the H^N^ saturation sequence^67^ that can be removed by its phase cycling and/or by appending a concatenated ^1^H spoil sequence (z-gradient 1 - 90°(^1^H) - z-gradient 2).

Residual dipolar couplings (RDCs) were determined from the difference between couplings observed for aligned versus unaligned (P3-7)_2_ samples (with U-^15^N,^13^C labeling) where alignment was achieved^68^ using a gel kit from New Era Enterprises, Inc. (Vineland, NJ, USA). For this, 7% acrylamide gels were dialyzed in ultrapure water for 3 h and NMR buffer overnight. The prepared gels were then soaked in the peptide sample (ca. 0.2 mM) overnight at 277 K and squeezed into open-ended 5 mm NMR tubes using a funnel and piston. The filled tube was closed with a bottom plug and a Shigemi top plunge. The sample alignment uniformity was assessed via the deuterium signal splitting. One bond ^1^H-^15^N RDCs were obtained from comparing aligned versus unaligned 2D BEST-TROSY spectra^59^ measured at 278 K and selecting either the H^N^ TROSY or semi-TROSY signals in the direct dimension. One bond ^13^C’-^15^N and two bond ^13^C’-^1^H^N^ RDCs were derived by comparing the pertaining ^1^J_C’,N_ splitting in the indirect (^15^N) and ^2^J_C’,H_ splitting in the direct (H^N^) dimension, respectively, observed in the (not ^13^C decoupled) 2D ^1^H,^15^N HSQC spectra of aligned *vs* unaligned samples measured at 278 K. PALES^69^ was used to calculate the expected RDCs for each structure in the CoMAND-derived ensemble, allowing for the calculation of an independent alignment tensor for each frame based only on its coordinates. Predicted RDCs were obtained as the average of the *ca.* 200 frames generated in the 20 iterations of converged, R-factor-minimizing CoMAND global ensemble calculations.

To study molecular recognition we measured the 2D ^1^H-^15^N BEST-TROSY spectrum of ^15^N-labeled Rap74-CTD (50 μM throughout, 298 K) in the presence of increasing concentrations of peptide. Before the experiment, both the protein and the peptide were dialyzed (277 K, two dialysis steps) in the same preparation of NMR buffer (20 mM sodium phosphate, 0.1% trifluoroacetic acid, pH 7.4), using a Pur-A-Lyzer (Sigma-Aldrich, Burlington, MA, USA) and a Micro Float-A-Lyzer (Spectrum Laboratories, San Francisco, CA, USA) respectively. The chemical shift assignment of RAP74-CTD was reported previously (BMRB code 27288). Averaged ^1^H and ^15^N chemical shift perturbations (CSPs) were calculated as:

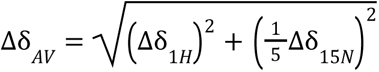

Dissociation constants, K_D_, and averaged CSP amplitude, Δδ_max_, for δ and γ peptide binding to Rap74-CTD were obtained by a global fitting (nonlinear regression) of the 10% peaks with the largest averaged CSPs to the following single-site binding model (1:1 stoichiometry):

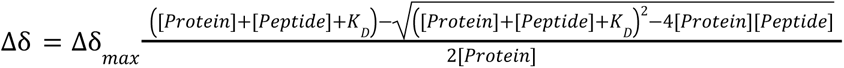

A Monte Carlo simulation varying both protein and peptide concentrations within 20% experimental errors was used to derive error margins for the final K_D_ values.

### Structural characterization using CoMAND

To investigate the conformational tendencies of the (P3-7)_2_ peptide we applied the CoMAND method (<u>Co</u>nformational <u>M</u>apping by <u>A</u>nalytical <u>N</u>OESY <u>D</u>ecomposition^27^). This method analyzes a 3D CNH-NOESY spectrum (i.e. 3D [H]C,NH HSQC-NOESY-HSQC), which displays only NOE contacts between ^15^N-bound and ^13^C -bound protons and is therefore intrinsically diagonal free^65^. As a first step, one-dimensional ^13^C sub-spectra (strips) were extracted from this spectrum (τ_m_=400 ms). Each strip is taken perpendicular to a specific ^15^N-HSQC position and represents contacts to a single ^15^N-bound proton, edited by the ^13^C shift of the attached carbon. For (P3-7)_2_ we obtained strips for 18 main chain amide protons (residues L7 to K24) and all 4 glutamine side chain H_ε21_ protons (Q11, Q14, Q17, Q20). These strips are analyzed in terms of a quantitative R-factor expressing the agreement between experimental and back-calculated spectra. Global back-calculation parameters for CoMAND were optimized by grid searching, resulting in an overall correlation time of 2.0 ns and effective ^13^C signal halfwidth of 14 Hz.

For a reconstruction of the experimental ^13^C strips, CoMAND compiles a linear combination of strips back-calculated from a set of trial conformers that should reflect the conformational space of each residue. Here, we used the a99sb-disp and DES-amber MD trajectories and back-calculated 13002 frames from each trajectory for each of the 22 experimental strips. For each residue, the conformational ensemble producing the lowest R-factor was then compiled using the CoMAND stochastic optimisation method^27^. A starting conformer is randomly selected from the 20 conformers with lowest R-factors. All conformers are then tested in random order, with a new member added to the ensemble if it decreases the R-factor by more than a given threshold (0.0005). Convergence is achieved if no further conformer is found or if the ensemble reaches a maximum size, here set at 20 structures. Due to its stochastic nature, this selection procedure can be repeated to produce ensembles with similar R-factors, but sampling a wider range of conformers.

For (P3-7)_2_ we applied a two-step protocol for each residue. In the first step we established the minimum R-factor by compiling 100 per-residue ensembles, optimizing over single experimental strips. These per-residue ensembles were also used to define a set of *witness* strips; i.e. those whose R-factors may be affected by conformational changes in the residue in question. In the second step, we obtained the conformational distribution by co-optimizing over these witness strips. For each set, 100 optimization trials were run for each MD trajectory frame pool, resulting in 100 ensembles per frame pool, each typically containing 5 to 15 members. After removing ensembles with R-factors significantly above average (90% confidence interval), the set of conformers used in co-optimization (typically over 1000) was pooled to represent the conformational diversity for each residue.

To quantify the conformational distributions, we clustered the data via Gaussian mixture modeling (GMM). A vector of *n* features - here dihedral angles - was defined for each conformer which was then used to train a model describing the probability p(k|x) that a data point x is a member of cluster k. For each cluster, this probability is defined by an *n*-dimensional multivariate Gaussian distribution representing its center and shape, and by a prior probability, p(k), corresponding to its relative population. These model parameters were fitted to the training data using the Expectation Maximization (EM) algorithm, modified to accommodate the periodic nature of dihedral angles. For (P3-7)_2_, we applied the GMM method for the χ_1_/χ_2_ pairs of all leucine and glutamine residues. As the number of clusters that best describe the training data was not known *a priori,* we searched values from 1 to 9 systematically and assessed the fit via the Bayesian information criterion, a measure that includes a penalty for model complexity. EM initialisation requires an arbitrary seed value for each cluster center. For χ_1_/χ_2_ pairs, it is convenient to select seeds at the center of a rotameric form, with seeds progressively added with increasing cluster number, according to their database frequency. The best scoring model was stored for each residue.

The GMM method provides a compact but detailed description of conformational landscapes for use in downstream calculations. Here we have applied it to Monte-Carlo conformational sampling as part of an extended “greedy” R-factor optimisation protocol. The pooled a99sb-disp and DES-amber MD trajectories were systematically sampled and the conformer affording the greatest reduction in global R-factor added in each iteration. Thus, 2-4 conformers were typically added to the ensemble, which was then further optimized by adjusting the side chain conformations for leucine and glutamine residues using χ_1_/χ_2_ combinations from the corresponding GMM model with a 0.05 probability cutoff. For each residue in the ensemble, up to 30 χ_1_/χ_2_ combinations were tested to find sterically acceptable conformations lowering the global R-factor. For glutamine, the χ_3_ angle was additionally sampled around population centers pertaining to each χ_1_/χ_2_ combination (5 trials; standard deviation 8°). Note that enthalpic contributions from hydrogen bonding were not considered in testing conformers, such that their selection was primarily driven by the reduction in R-factors. This iterative process of ensemble selection and modification was repeated until no further conformers were added by the greedy step. This protocol was repeated 20 times to probe the consistency of results and an example chosen as the final ensemble (Fig. 2g).

### Molecular Dynamics simulations

The trajectories for the (P3-7)_2_ peptide used in the CoMAND analysis and for the TBP peptide shown in Fig. 5h were generated with the Well Tempered Ensemble (WTE)^39,70^ enhanced sampling algorithm. We used 26 energy biased replicas within the temperature range from 275 K to 500 K, and two unbiased replicas at 278 K and 298 K. The unbiased replica at 278 K (at which temperature the 3D [H]C,NH HSQC-NOESY-HSQC was measured) was then used to generate the conformations for our CoMAND analysis and Fig 5h. For the biased replicas, the energy bias was increased during the first 500 ps and then kept constant. All replicas were subsequently used in a production simulation for 200 ns, where conformations used by CoMAND were extracted from the last 150 ns. Exchange between replicas was monitored to ensure good replica diffusion in temperature space. The production simulation was run in the NPT ensemble. We used two recent force fields for these simulations, DES-amber^28^ and a99sb-disp^29^, each with its pertaining TIP4P-D water model. Bonds with hydrogen atoms were constrained and a time-step of 2 fs was used. For our simulations we used Gromacs 2019.4^71–73^ patched with the PLUMED library^74^ version 2.5.3^75^ to enable the WTE sampling method.

Two sets of trajectories for segments of L_3_X peptides (see Table S1) with the sequence L^1^L^2^L^3^XQ^1^Q^2^Q^3^Q^4^Q^5^Q^6^Q^7^Q^8^ (where X is any of the 20 natural proteinogenic amino acids) were calculated using either the Charmm36m force field^33^ with a TIP3P^76^ water model or the a99sb-disp force field^29^ with its TIP4P-D water model. To bias the simulations towards relevant conformations, calculations were started from a fully helical conformation and the backbone φ and ψ dihedral angles of residues L^2^ to Q^5^ were restrained to -60° and -40°, respectively, with a spring constant *k* value of 5 kJ mol^-1^ degree^-2^. Similarly, a bias to increase the occurrence of Q^4^ → X side chain to main chain hydrogen bonds was introduced by restraining the distance between the main chain O of residue X and the N_ε2_ of residue Q^4^ to 4 Å, with a spring constant *k* value of 50 kJ mol^-1^ nm^-2^. Structure minimization and thermalization at 278 K was performed in the NVT ensemble for 1 ns. For the 1 μs NPT production runs we used Gromacs 2019.4^71–73^ and an equilibration period of 100 ns that was excluded from trajectory analysis.

A trajectory was calculated for peptide Q1E using the CHARMM22*^77^ force field and TIP3P water model^76^ to obtain different structures with i-4 → i side chain to main chain hydrogen bonds involving the glutamic acid carboxyl group that served as seeds for the QM/MM simulations (see below). The dihedral angle χ_4_ orienting the glutamic acid carboxyl group was restrained to 0° since its most stable conformation in CHARMM22*, corresponding to χ_4_ = 180°, is incompatible with the experimentally indicated side chain to main chain interaction. A fully helical starting structure was thermalized (300K) and equilibrated in the NVT ensemble for 1 ns. The 1 μs production run was obtained using ACEMD^78^ with a 100 ns equilibration period.

### QM/MM simulations

Four starting structures were selected from the classical MD trajectory of peptide Q1E conserving their box of water and ions. For the QM/MM simulations we used AMBER 20^79^ coupled to the QM Terachem 1.9 interface^80–82^. The QM subsystem was described at the BLYP/6-31G* level of theory including dispersion corrections^83^ and comprised 66 atoms including linker atoms. The classical subsystem was treated with the CHARMM22*^77^ and TIP3P^76^ force fields. The linker atom procedure was employed to saturate the valency of the frontier atoms and electrostatic embedding was used as implemented in AMBER. An electrostatic cutoff of 12 Å and periodic boundary conditions were employed throughout all QM/MM MD simulations, using a time step of 1 fs. Structures were minimized, thermalized, and equilibrated for 10 ps at the QM/MM level prior to the 150 ps-long production runs. Finally, for each of the 150 ps QM/MM-MD runs, a Natural Bond Critical Point analysis^84,85^ was performed using NBO 7.0^86^.

### Database motif searches

UniprotKB^34^ (including both the Swiss-Prot and the TrEMBL databases) was queried for protein sequences containing motifs that fulfill our design rules using the ScanProsite^35^ motif search tool hosted at the Expasy website (https://prosite.expasy.org/scanprosite/). The query motif was introduced in Prosite format as: [LFYWIM]-X-X-[LFYWIM]-Q-X-[LFYWIM]-Q-X-[LFYWIM]-Q-X-Q-X-X-Q-X-X to quest for sequences with n = 4 Q_i+4_ - [LFYWIM]_i_ pairs, with the number of central -[LFYWIM]-Q-X- triplets increased stepwise for concomitant increases in the quested number of Q_i+4_ - [LFYWIM]_i_ pairs. UniprotKB annotation scores and taxonomic lineage information was obtained by programmatically accessing this information in UniprotKB using the accession codes from the ScanProsite searches. The same codes were used to retrieve structural models from the AlphaFold Database^18^, in which the secondary structure of the (P3-7)_n_-like motifs fulfilling our design rules was obtained using DSSP^36^.

### Hydrogen bond criteria

We considered two atoms to be hydrogen bonded if the distance between donor H and the acceptor O was shorter than 2.4 Å and their angle was larger than 120°.

## Acknowledgements

We thank Borja Mateos, Ben Lehner, Ernest Giralt and Luis Serrano for helpful discussions and the ICTS NMR facility, managed by the scientific and technological centers of the University of Barcelona (CCiT UB), for their help in NMR. X.S. acknowledges funding from AGAUR (2017 SGR 324), MINECO (BIO2015-70092-R and PID2019-110198RB-I00), and the European Research Council (CONCERT, contract number 648201). B.B.K acknowledges funding from the Novo Nordisk Foundation (#NNF18OC0033926). MO acknowledges funding from the Instituto Nacional de Bioinformática, The EU BioExcel Centre of Excellence for HPC and the Spanish Ministry of Science (RTI2018-096704-B-100) and the Instituto de Salud Carlos III–Instituto Nacional de Bioinformatica (ISCIII PT 17/0009/0007 co-funded by the Fondo Europeo de Desarrollo Regional). MO is an ICREA Academy scholar and JA is a Juan de la Cierva fellow. IRB Barcelona is the recipient of a Severo Ochoa Award of Excellence from MINECO (Government of Spain).

## Supplementary Figures

**Fig. S1.**
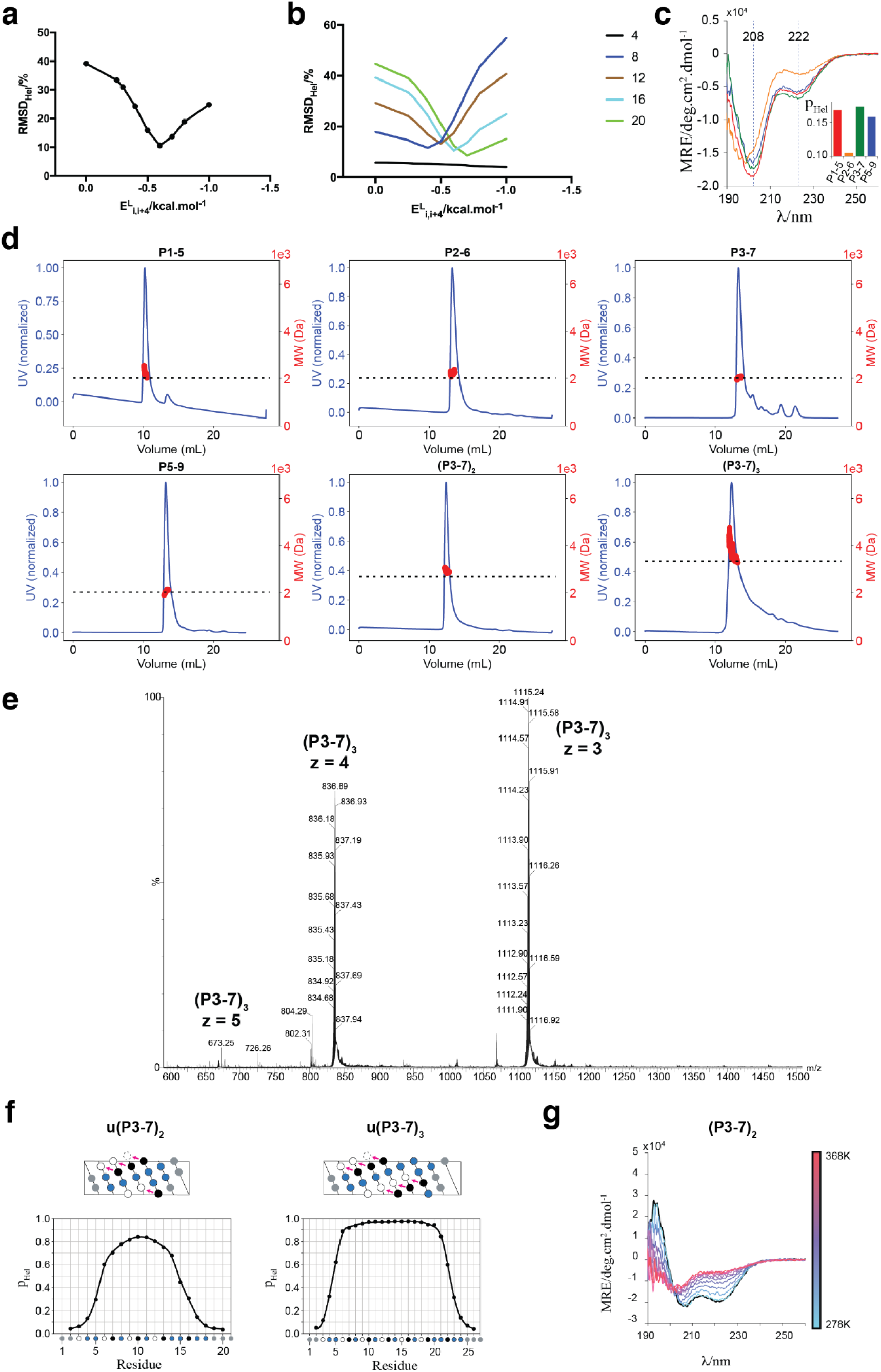
Cooperativity of Gln_i+4_→Leu_i_ bifurcated hydrogen bonds in monomeric polyQ tracts and designed sequences. **a** Optimizing the value of a newly introduced energy term E^L^i,i+4 in *Agadir* to account for the Gln_i+4_➝Lei_i_ interaction improves its accuracy at predicting the helicity of L_4_Q_16_ (Table S1). **b** The optimal value of E^L^i,i+4 depends on polyQ tract length. Numbers in the legend indicate the length of the L_4_Q_n_ peptides (full sequence KKPGASL_4_Q_n_KKY). **c** Overlay of the CD spectra obtained for the peptides in Fig. 1a. The dashed vertical lines highlight the positions where minima are observed in the CD spectrum of the α-helix (208 and 222 nm). In the inset, average peptide helicities are shown as obtained from deconvoluting the CD curve using the algorithm CONTIN (reference set 7) hosted at DichroWeb (http://dichroweb.cryst.bbk.ac.uk). **d** SEC-MALS analyses of the peptides in Fig. 1a and 1b. The horizontal dashed line indicates the molecular weight of the monomeric species calculated using the Protparam algorithm hosted at Expasy (https://web.expasy.org/protparam/). **e** Native mass spectrum of a ^13^C-^15^N labeled sample of the peptide (P3-7)_3_, as obtained by Q-TOF MS using a Synapt G1-HDMS (see supplementary methods). Three ionization states corresponding to the monomeric species were independently detected. No oligomeric species were detected. **f** Chemical shifts-derived helical propensity (p_Hel_) profiles of the u(P3-7)_2_ and u(P3-7)_3_ peptides, which are versions of the peptides shown in Fig. 1b lacking the PGAS N-capping sequence (the ‘*u*’ stands for *uncapped*). **g** Thermal unfolding of (P3-7)_2_. Shown are CD spectra acquired every 10 K in the 278 K - 368 K range (blue to red color gradient). The CD spectrum back at 278 K after thermal unfolding is shown in black.

**Fig. S2.**
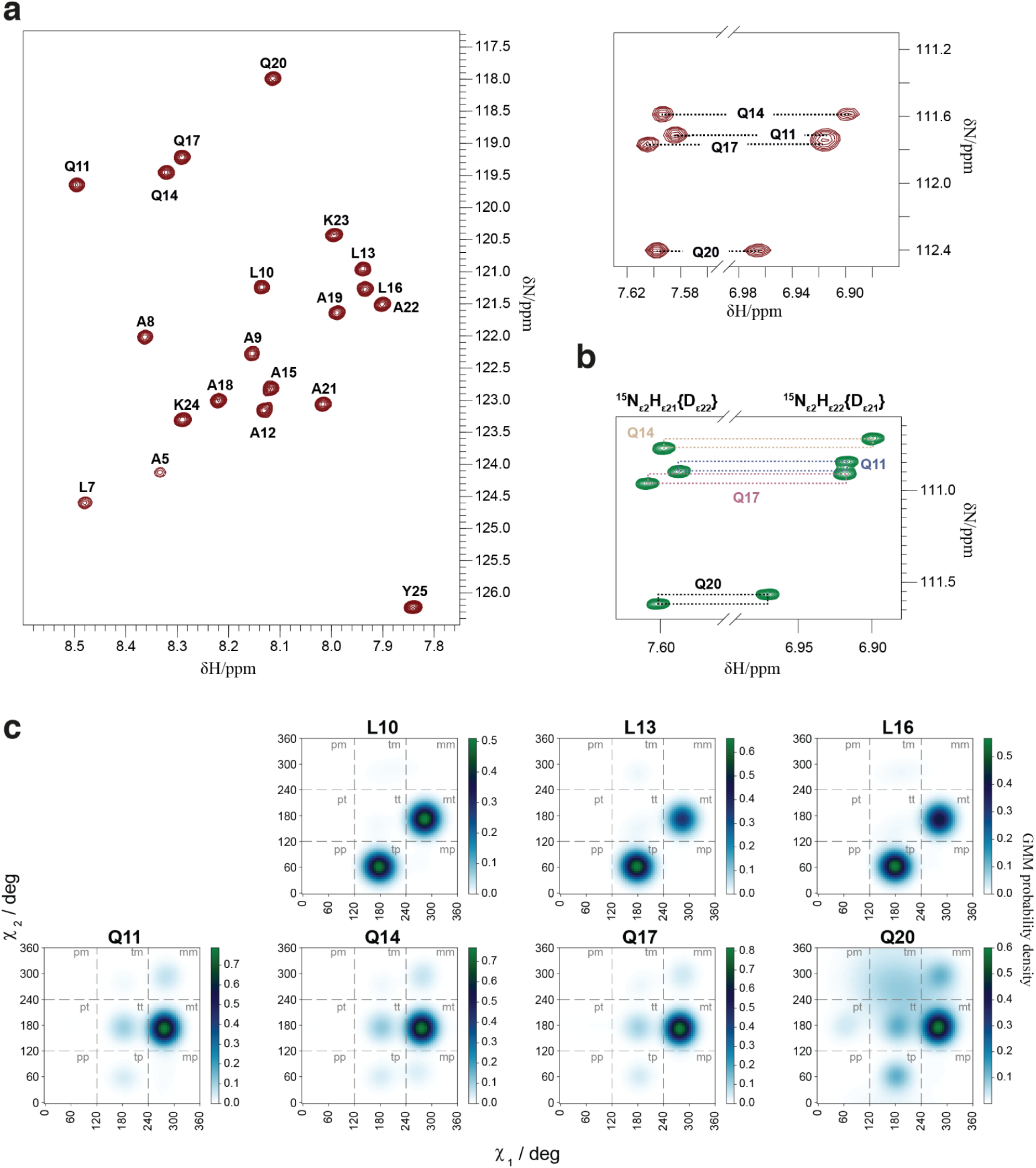
Structural features of Gln_i+4_➝Leu_i_ bifurcated hydrogen bonds: supporting analyses of the peptide (P3-7)_2_. **a** Assigned ^1^H-^15^N HSQC spectrum of the peptide (P3-7)_2_. Left: region of the spectrum displaying main chain N-H correlations. The spectrum shows wide signal dispersion in both dimensions and no signal overlap, which was exploited for the structural characterization shown in Fig. 2. Right: region of the spectrum showing the Q side chain NH_2_ correlations. **b** Side chain region of a ^1^H-^15^N HSQC spectrum acquired on a (P3-7)_2_ sample in 50% D_2_O applying deuterium decoupling during the ^15^N evolution time. The differential deuterium isotope shift in the NHD species results in different ^15^N chemical shifts for each of them, which allows the unambiguous stereospecific assignment of the side chain carboxamide H_ε_ resonances^66^ **c** Probability densities for χ_1_ and χ_2_ of all residues captured by the Gaussian Mixture Model (GMM) from the CoMAND per-residue fits.

**Fig. S3.**
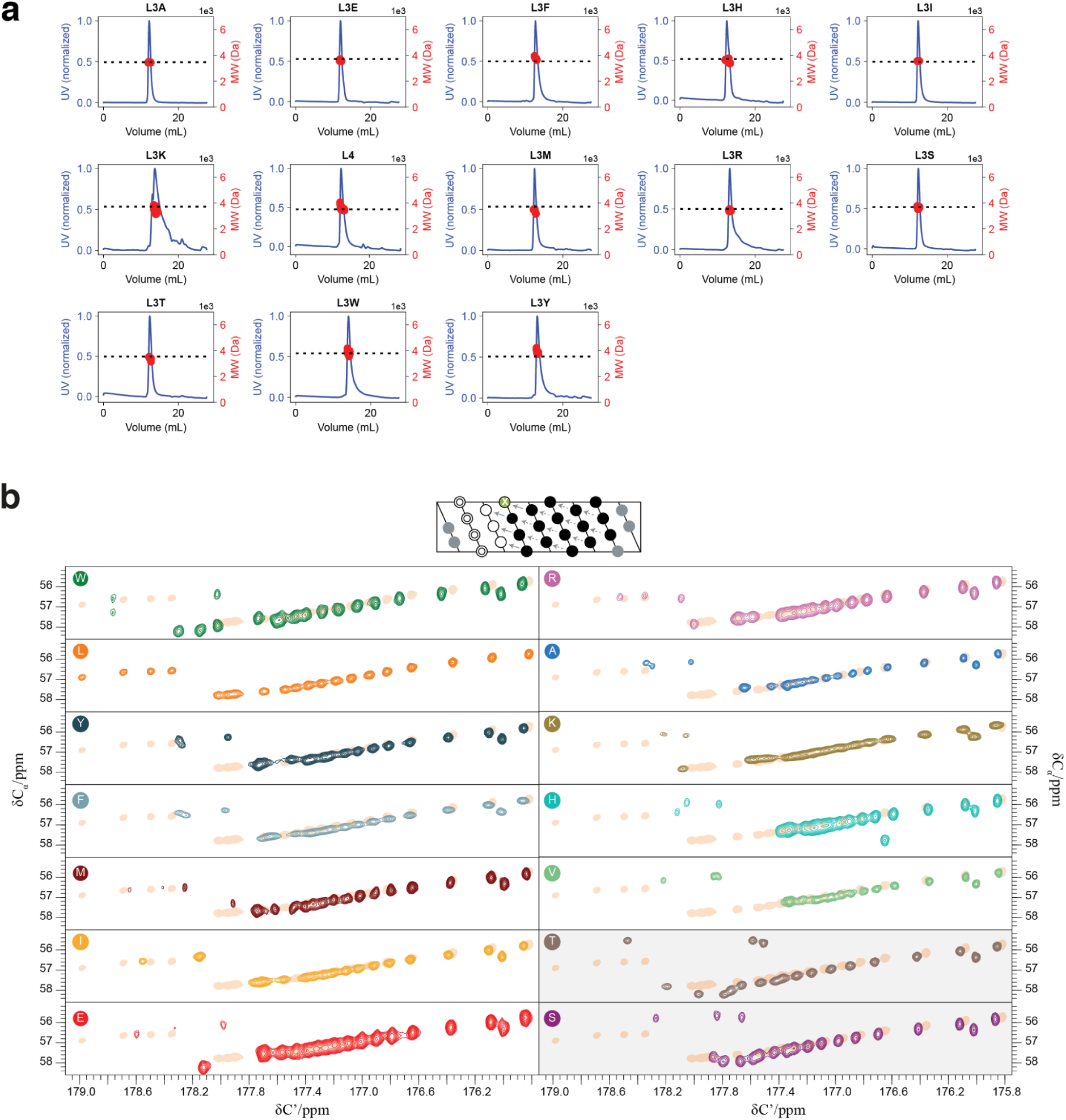
SEC-MALS analyses and CACO spectra of the L_3_XQ_16_ peptides. **a** SEC-MALS analyses of the peptides in the L_3_XQ_16_ series shown in Fig. 3. The horizontal dashed line indicates the molecular weight of the monomeric species calculated using the Protparam algorithm hosted at Expasy (https://web.expasy.org/protparam/). **b** ^13^C-detected CACO spectra of the L_3_XQ_16_ variants, sorted by their observed helicity, with the spectrum of L_4_Q_16_ underlaid (orange shadow) in every panel for reference. Spectra shaded in gray indicate variants with outlier behavior.

**Fig. S4.**
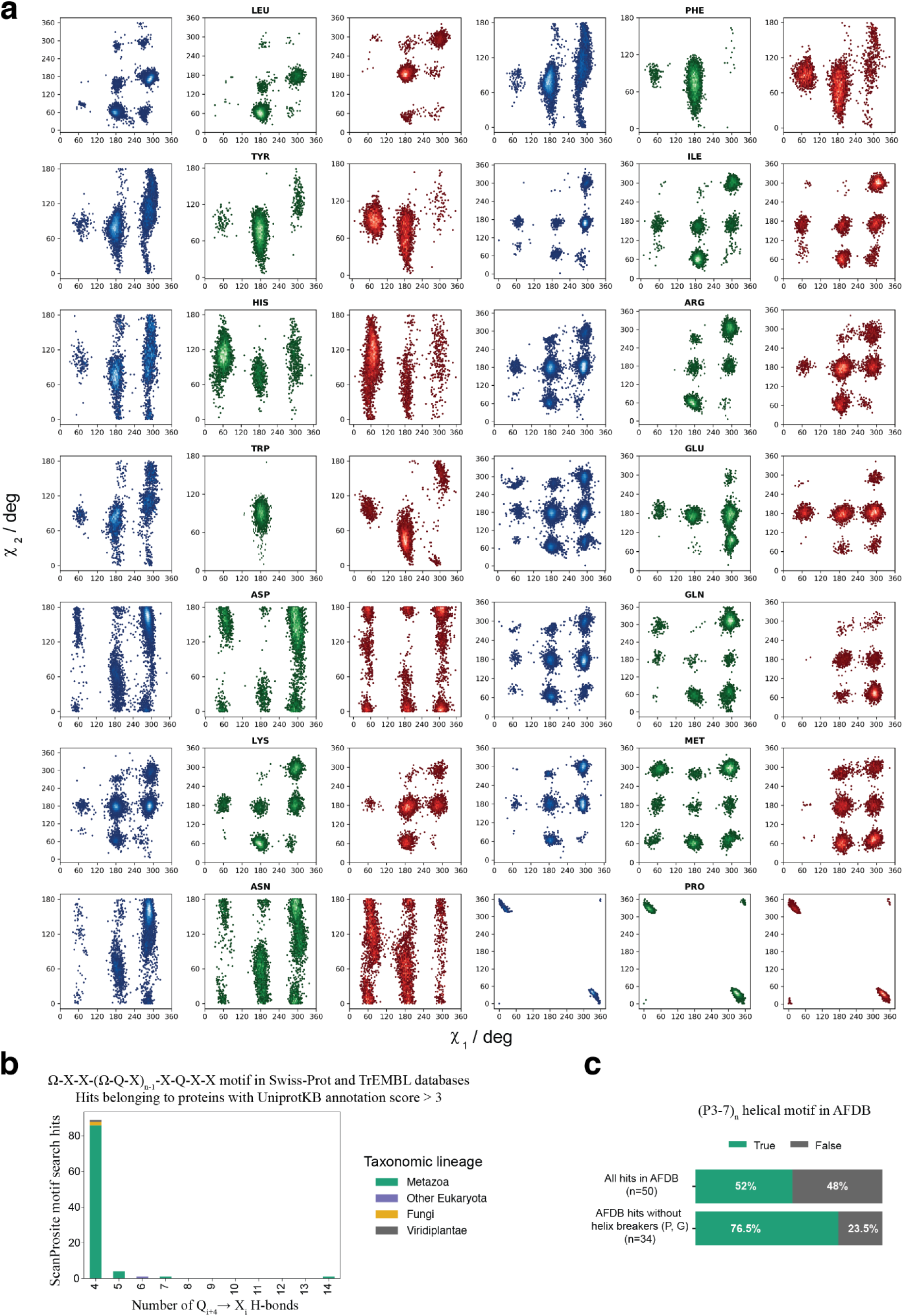
Rotamer sampling in the L_3_XQ_16_ MD simulations and supporting data on the natural sequences fulfilling our design rules. **a** Side chain χ_1_-χ_2_ dihedral angle correlations for residues with at least three side chain carbon atoms. In blue, values as reported in the BBRep database for residues located in the context of a helix. In green, values sampled throughout the Charmm36m 1 μs MD trajectories calculated for the corresponding L_3_XQ_16_ variants. In red, values sampled throughout the a99sb-disp 1 μs MD trajectories calculated for the corresponding L_3_XQ_16_ variants. **b** Number of ScanProsite-identified^35^ protein sequences with an UniprotKB^34^ annotation score > 3 found in the Swiss-Prot and TrEMBL databases containing the (P3-7)_n_ motif (Ω = W, L, Y, F, I and M; X = any amino acid). Counts are grouped by taxonomic lineage as obtained from UniProt. **c** 50 proteins containing sequence motifs fulfilling our design rules have models in the AlphaFold Database (AFDB)^18^. The top bar shows the percentage of these proteins where the (P3-7)_n_-like motif is predicted to be helical from residue 1 to residue -4, as obtained from a DSSP analysis^36^ of the structural models. The bottom bar shows the same for proteins not containing helix breaking residues (P, G) in the same segment (n=34).

**Fig. S5.**
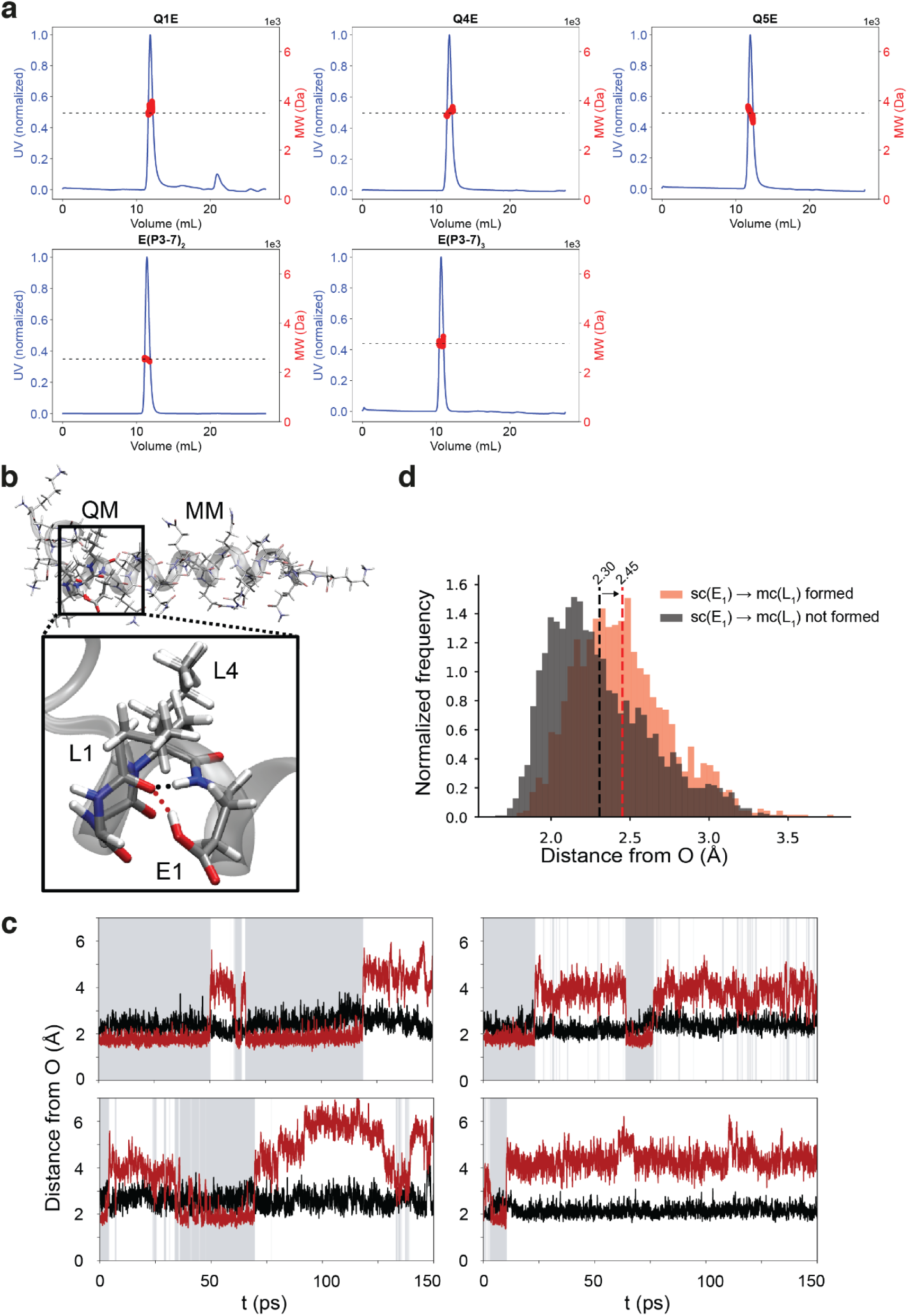
A pH-sensitive conformational switch based on glutamic acid bifurcated hydrogen bonds. **a** SEC-MALS analyses of the QXE and E(P3-7)_n_ peptides shown in Fig. 4.The horizontal dashed line indicates the molecular weight of the monomeric species calculated using the Protparam algorithm hosted at Expasy (https://web.expasy.org/protparam/). **b** Schematic representation of the QM/MM calculations setup. The atoms included in the QM subsystem are shown in sticks and the distances in **c** are indicated with dashed lines. **c** Time series of hydrogen bond donor-acceptor distances throughout four independent QM/MM trajectories. In black, H^N^-O distance. In maroon, H_ε2_-O distance. Frames in which bifurcated hydrogen bonds are observed are indicated shadowed in gray. **d** Distribution of the distance between the main chain NH of E1 and the main chain CO of L1 in the absence and in the presence of the sc(E1)→mc(L1) hydrogen bond.

**Fig. S6.**
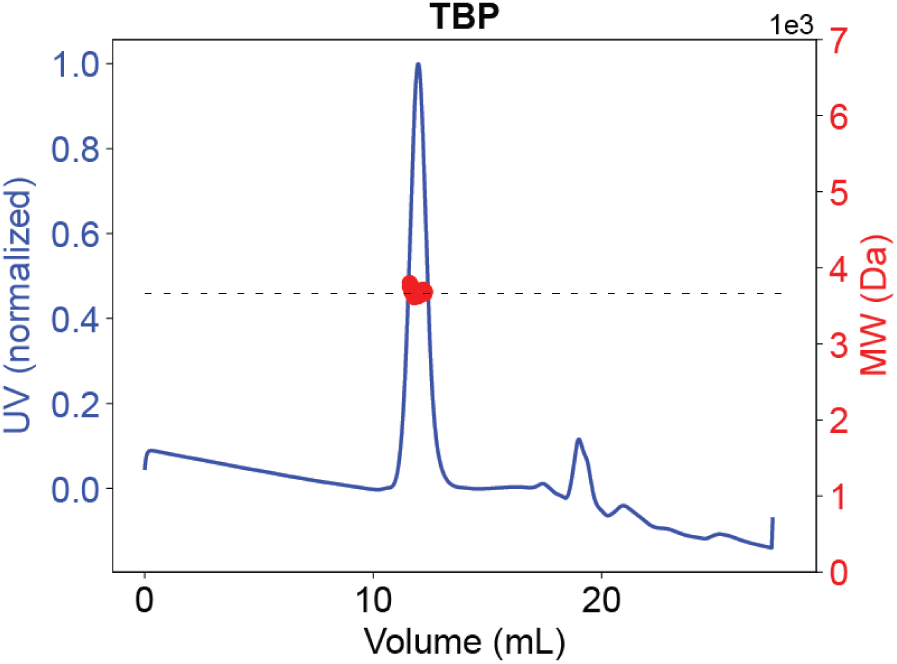
SEC-MALS analysis of the TBP polyQ peptide. Supporting SEC-MALS analysis for the peptide shown in Fig. 5. The horizontal dashed line indicates the molecular weight of the monomeric species calculated using the Protparam algorithm hosted at Expasy (https://web.expasy.org/protparam/).

**Fig. S7.**
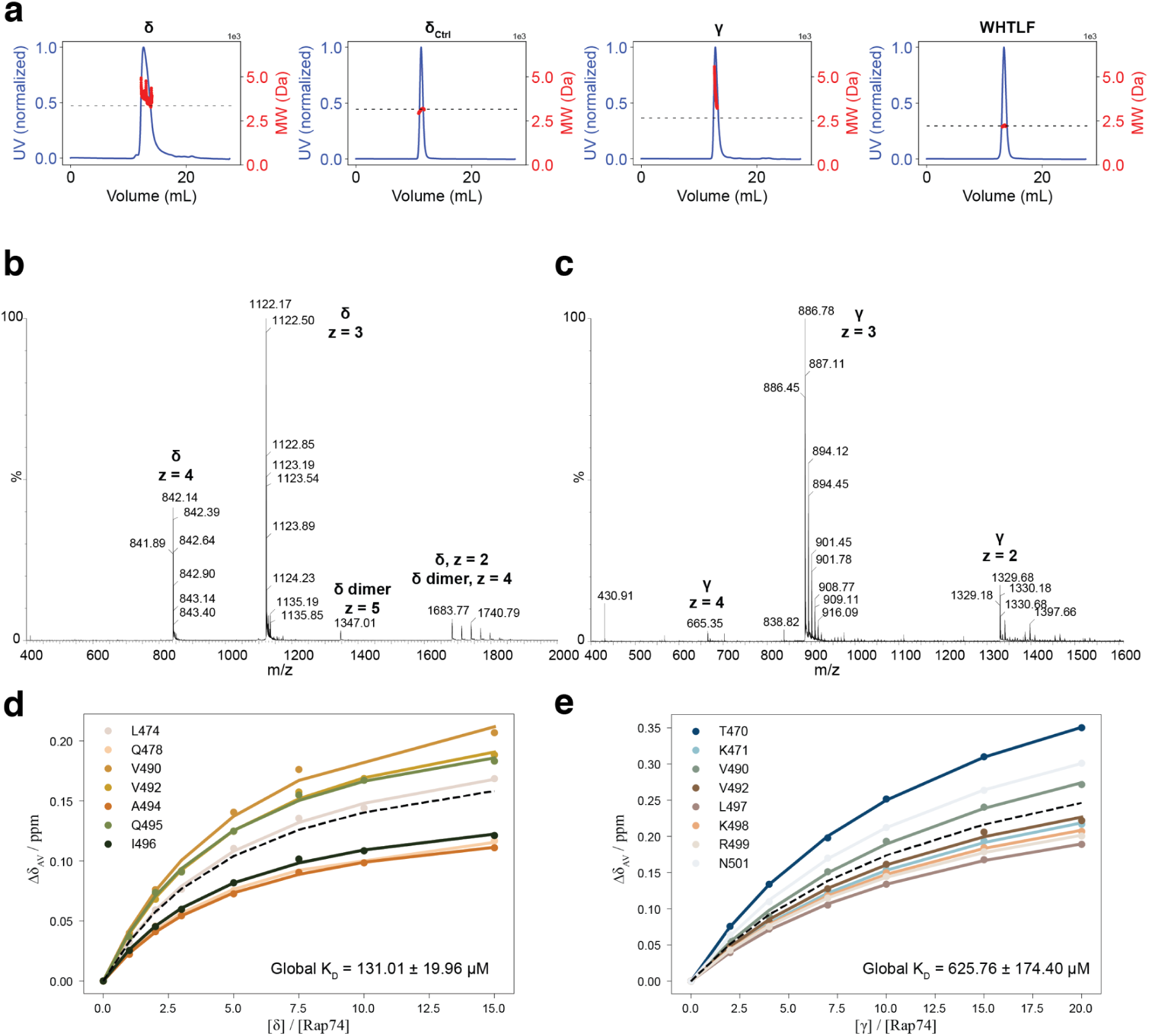
Design of binding motif-grafted peptides to bind the Rap74 domain of TFIIF: supporting analyses. **a** SEC-MALS analyses of the peptides shown in Fig. 6. The horizontal dashed line indicates the molecular weight of the monomeric species calculated using the Protparam algorithm hosted at Expasy (https://web.expasy.org/protparam/). **b** Native mass spectra of the peptide δ as obtained by Q-TOF MS using a Synapt G1-HDMS (see supplementary methods). Three ionization states corresponding to the monomeric species were independently detected. Two ionization states (one resolved, one ambiguous) corresponding to traces of the dimeric species were detected. **c** Native mass spectra of the peptide γ. Three ionization states corresponding to the monomeric species were independently detected. No oligomeric species were detected. **d** K_D_ for the binding of the δ peptide to Rap74 as obtained from a global fit (black dashed line) of the binding curves for the top 10% peaks with most intense averaged CSPs (individual fits shown as solid color lines). **e** As in **d** for the binding of the γ peptide to Rap74.

**Table S1.**
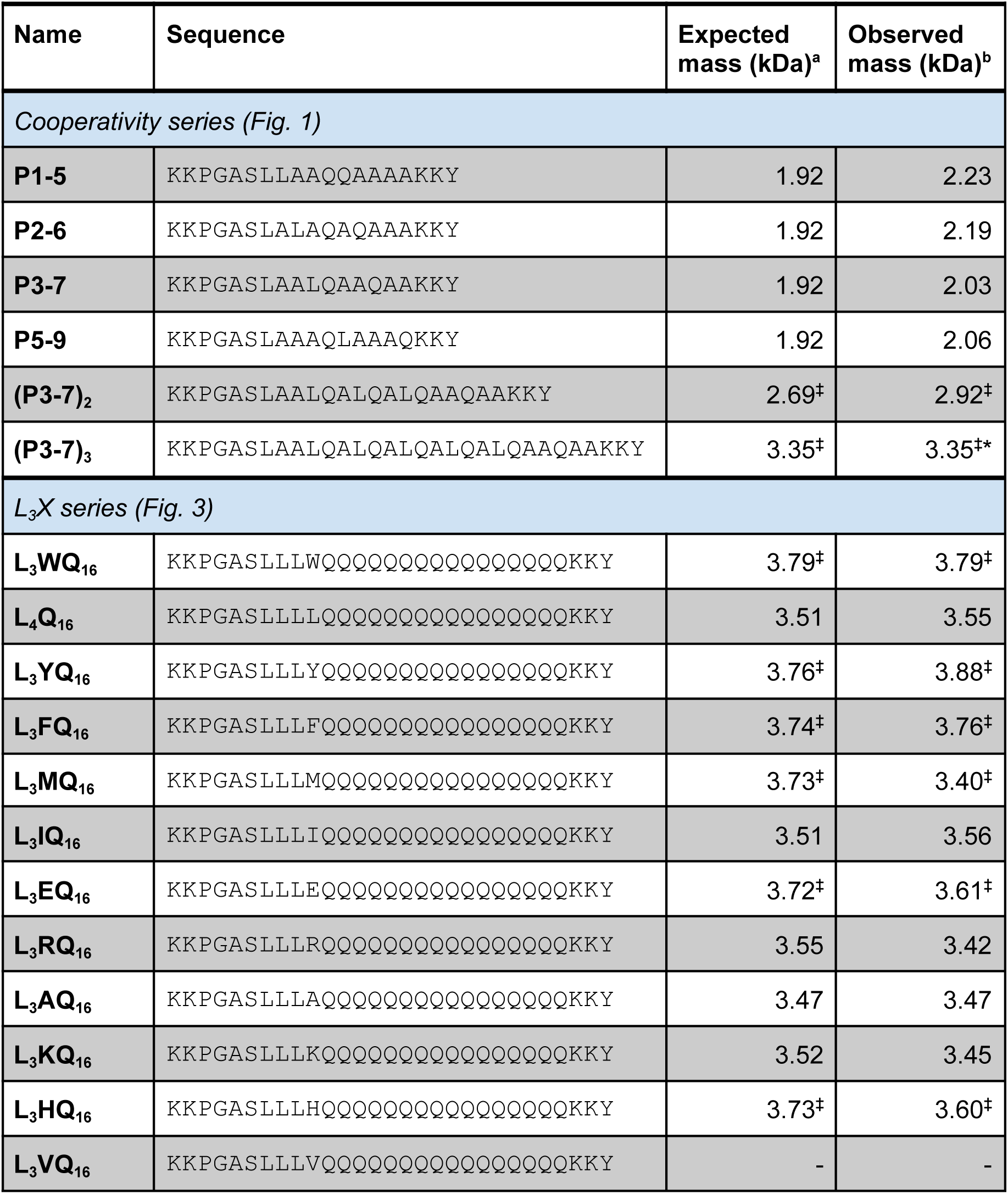
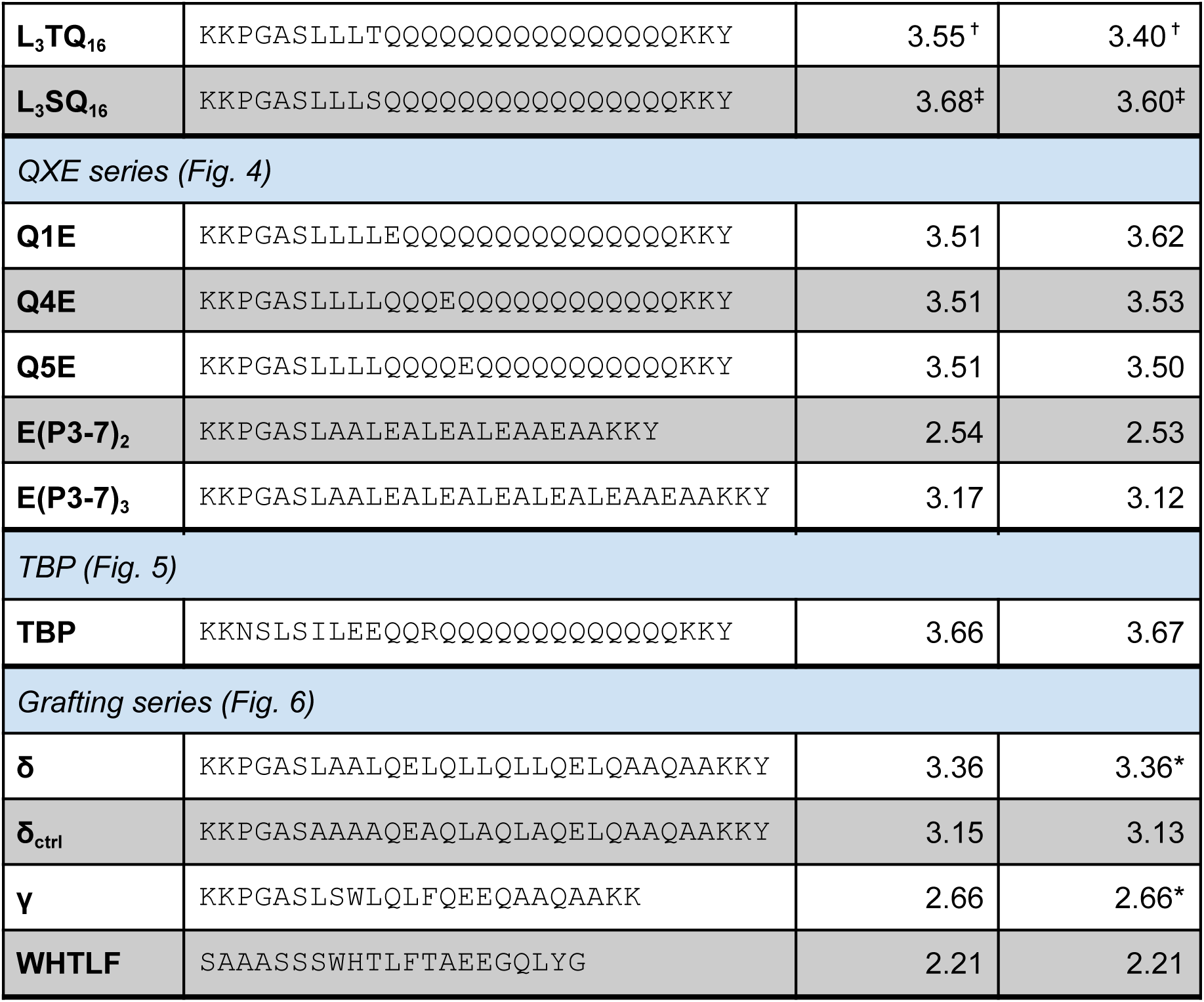
Details of the peptides used in this study. **a** No accompanying symbol indicates that the mass reported was calculated considering ^12^C and ^14^N isotopes using the Protparam algorithm hosted at Expasy (https://web.expasy.org/protparam/). ✝ indicates the mass calculated for the ^15^N enriched species and ‡ indicates the mass calculated for the ^13^C-^15^N enriched species. **b** Isotope labeling scheme as in **a**. In addition, no accompanying symbol indicates that the reported mass is an average of that measured during the corresponding peak elution using SEC-MALS, whereas * indicates that the reported mass is a z = 1 deconvoluted average of the main peaks corresponding to all detected ionization states in the native MS spectra.

## Supplementary Methods

### Parametrization of Q_(i+4)_ → L_(i)_ side chain to main chain hydrogen bonds in Agadir

Agadir source code was kindly shared by Professor Luis Serrano (Center for Genomic Regulation, Barcelona, Spain). The energy term accounting for Leu_i_ - Gln_i+4_ interactions E^L^_i,i+4_ was progressively increased in the 0 to 1 kcal mol^-1^ range. Chemical shift-derived (BMRB entries 27713, 27714, 27715, 27716, 27717) experimental per-residue helical propensity (p_hel_) values for the L_4_Q_n_ peptides (full sequence KKPGASL_4_Q_n_KK) were previously reported^11^. At each E^L^_i,i+4_ increase step, agadir was used to predict per-residue helicity and the RMSD between the predicted and experimentally determined per-residue helical content was obtained using:

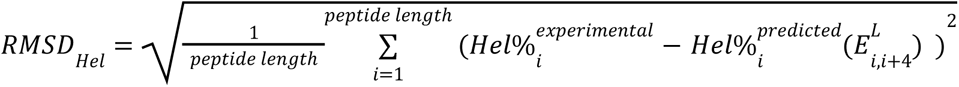

### SEC-MALS analyses

The oligomeric state of peptides in solution was determined by size exclusion chromatography coupled to multiple angle light scattering. Peptide samples were loaded in a Superdex Peptide 10/300 GL column (GE Healthcare, Chicago, IL, USA) mounted on a Shimadzu Prominence Modular HPLC with a SPD-20 UV detector (Shimadzu, Kyoto, Japan) coupled to a Dawn Heleos-II multi-angle light scattering detector (18 angles, 658 nm laser beam) and an Optilab T-rEX refractometer (Wyatt Technology, Santa Barbara, CA, USA). The SEC-UV/MALS/RI system was equilibrated with sodium phosphate 20 mM, 0.1% TFA at 298 K with a 0.5 mL min^-1^ flow rate and elution was monitored using UV absorbance at 280 nm over 55 minutes. Data acquisition and processing was performed using the Astra 6.1 software (Wyatt Technology, Santa Barbara, CA, USA).

### Native Mass Spectrometry

Native mass spectrometry was used to determine the oligomeric state of some peptides. MS experiments were performed using a Synapt G1-HDMS mass spectrometer (Waters, Manchester, UK). All samples were buffer exchanged into 150mM ammonium acetate and were infused by automated chip-based nanoelectrospray using a Triversa Nanomate system (Advion BioSciences, Ithaca, NY, USA) as the interface. The ionization was performed in positive mode using a spray voltage and a gas pressure of 1.75kV and 0.5psi, respectively. The source pumping speed in the backing region (5.85 mbar) of the mass spectrometer was reduced to achieve optimal transmission of non-covalent complexes. Cone voltage, extraction cone and source temperature were set to 40V, 6V and 40°C, respectively. Trap and transfer collision energies were set to 6V and 4V, respectively. The pressure in the Trap and Transfer T-Wave regions were 2.39·10^-2^ mbar of Ar and the pressure in the IMS T-Wave was 0.467 mbar of N_2_. Trap gas and IMS gas flows were 8 and 25 mL/sec, respectively. The instrument was calibrated over the 300-8000Da m/z range using a solution of cesium iodide. MassLynx version 4.1 SCN 704 was used for data processing.

